# Evaluating the pathogenicity, stability, and stereochemical effects of variants of unknown significance in TREM2 using orthogonal structural analyses

**DOI:** 10.1101/2025.05.25.656054

**Authors:** Joshua Pillai, Kijung Sung, Linda Shi, Chengbiao Wu

**Affiliations:** School of Biological Sciences, University of California, San Diego, 9301 S Scholars Dr, La Jolla, CA 92093, USA; Department of Neurosciences, University of California San Diego, Medical Teaching Facility, 9500 Gilman Drive, La Jolla, CA 92093-0624, USA; Biophotonics Technology Center, Institute of Engineering in Medicine, University of California, San Diego, 9500 Gilman Dr, La Jolla, CA 92093, USA

**Keywords:** AlphaFold2, Alzheimer’s disease, TREM2, Neurodegenerative Diseases

## Abstract

Triggering receptor expressed on myeloid cells 2 (TREM2), an immune receptor expressed on the surface of microglia, has been identified through genome-wide association studies to be one of the risk factors in Alzheimer’s disease (AD). Several studies have also identified missense variants of TREM2 to be associated with frontotemporal dementia and Nasu-Hakola disease (NHD). To date, 51 novel missense variants of TREM2 have been identified in the literature, with the disease risk profiles of most variants still unknown. Assessing and classifying the pathogenicity of these variants is essential to investigate the disease mechanisms and develop effective treatments. Herein, we classified 32 missense variants involved in TREM2 using structural bioinformatic data with AlphaFold2. Using the protocol described in our previous work (Pillai et al., 2025), we determined the structural, stability, and potential functional effects of these variants. Our evaluations of the mutations were divided into those (i) implicated in NHD, (ii) located on the transmembrane domain, (iii) surface of the IgV-like domain, and (iii) buried in the receptor. Our analysis of variants involved in NHD suggests that, while V126G imposes the greatest effects, the T66M variant exerts significantly less effects on the TREM2 structures compared to the other remaining variants. Variants in the transmembrane domain of TREM2 did not impose significant alterations to the three-dimensional structure. Outside of known variants in the IgV-like domain, we identified 10 variants that imposed significant destabilizing effects to the structure and are of potential interest. Overall, the baseline biochemical data provided from this study may be informative to experimental efforts to better classify rare coding variants of TREM2 that are of unknown biological and clinical significance.

**Highlights:** Using our previously published protocol, we characterized the structure, stability, and functional effects of 27 unique variants of unknown biological significance in TREM2.

Our analysis was divided into variants located on the transmembrane domain, surface of the IgV-like domain, buried in the IgV-like domain, and those leading to Nasu-Hakola disease.

Excluding AD- and NHD-causing variants, we identified 10 TREM2 variants that cause significant destabilizing effects to the receptor.

The in-silico data provided from this study serve as a baseline, and can guide future experimental efforts involving TREM2 variants.

**Graphical Abstract:** 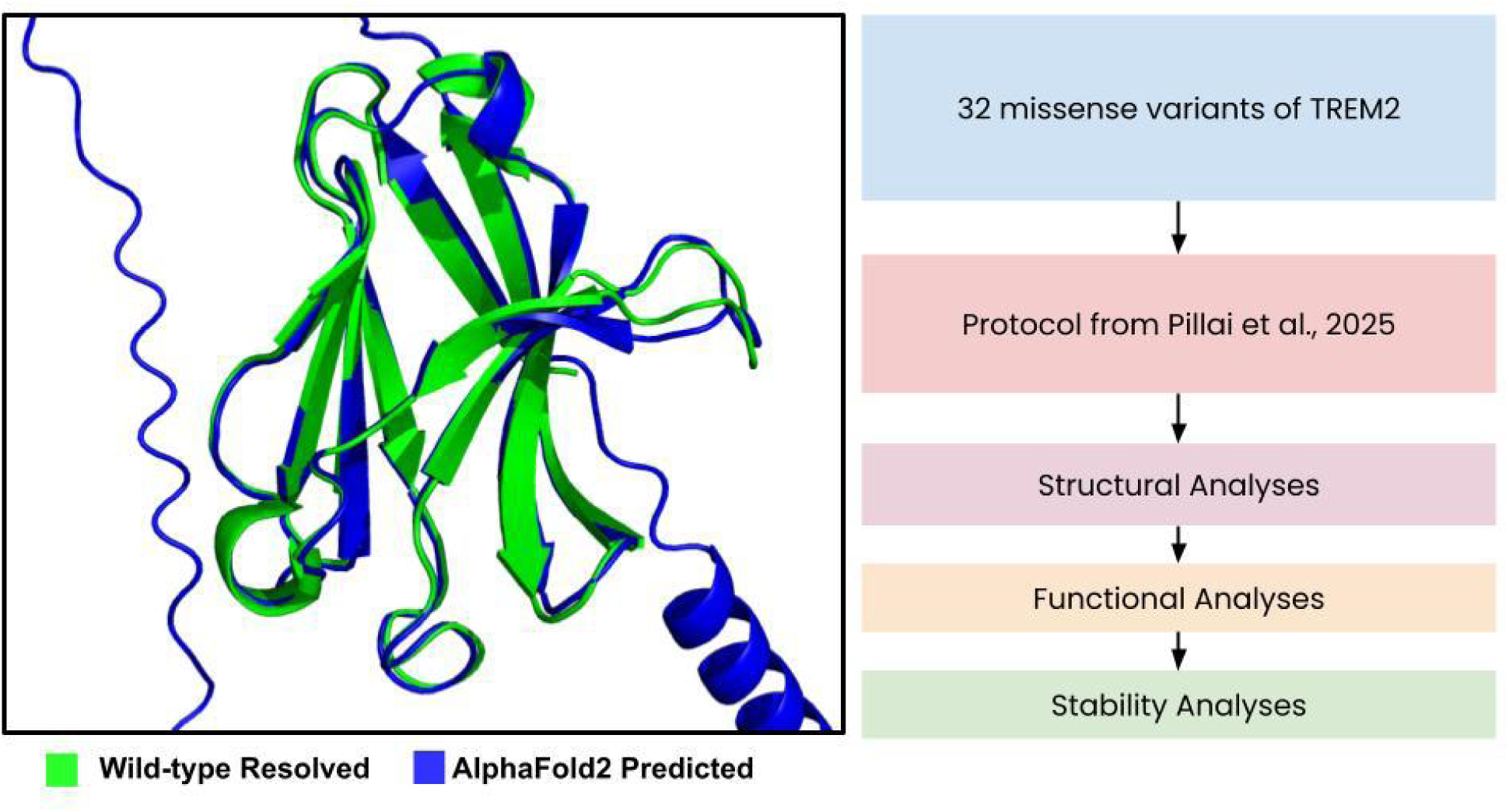

## 1. Introduction

Triggering receptor expressed on myeloid cells 2 (TREM2) is an immune receptor that is exclusively expressed on the surface of microglia found in the central nervous system (Li et al., 2023), and its variants have been implicated in Alzheimer’s disease (AD), frontotemporal dementia (FTD), and Nasu-Hakola disease (NHD) (Li et al., 2022; Deczkowska et al., 2020; Kulkarni et al., 2021). Notably, the R47H variant, identified by genome-wide association studies as a significant risk factor of Alzheimer’s disease, has been a focus of experimental efforts over the past decade (Jonsson et al., 2012; Guerreiro et al., 2012; Benitez et al., 2013; Ghani et al., 2016; Hooli et al., 2014; Sayed et al., 2021). Although the potential risk of R62H, D87N, and H157Y variants of TERM 2 has also been evaluated by meta-analysis, not enough evidence exists to confirm their relation to AD in comparison to those of R47H TREM2 (Li et al., 2021). Similarly to AD, the Y38C, W50C, T66M, and V126G variants have been found and validated in numerous studies as causal factors for NHD (Dash et al., 2020; Dardiotis et al., 2017; Kiianitsa et al., 2024; Bianchin et al., 2013; Carmona et al., 2017). Despite enormous experimental efforts have been taken to evaluate these variants, no systematic evaluation has been conducted on rare coding-variants of TREM2, and their potential clinical and biological effects.

In our previous report, we used AlphaFold2 (AF2) to obtain a high-confidence prediction of the complete TREM2 structure, and found that it was nearly identical to partially resolved structures from literature (Pillai et al., 2025). We introduced a systematic protocol to evaluate the structural, functional, and stability effects of missense mutations on unresolved single protein chains, and performed a case study with TREM2 R47H and validated known experimental evidence of this influential variant (Pillai et al., 2025). This encouraged further evaluations of the R62H and D87N with comparison of the Y38C and V126G variants, where we provided baseline predictions of their molecular details (Pillai et al., 2025). We believe that the in-silico data provided from the protocol may be informative and can serve as a validation of experimental efforts involved for this receptor. Herein, we used our previously described method on all known coding-variants of TREM2 that are located on high-confidence regions of the AF2-predicted three-dimensional structure. Out of 51 missense mutations known within current literature, we evaluated 32 variants that are located on high-confidence residues of the AF2 structure. We divided our analysis of the variants into those implicated in NHD, located on the transmembrane domain, surface of the IgV-like domain, and buried in the IgV-like domain. We believe that our data will aid experimental efforts to better define the pathogenicity of these rare variants.

## 2. Materials & Methods

### 2.1. 3D Structures

We used the AF2 structural model of TREM2 to determine the orthogonal effects of missense mutations on the 3D structure. From our prior study (Pillai et al., 2025), surprisingly the prediction from AF2 had the lowest root mean-square deviation (r.m.s.d.) to known WT structures derived from Sodom et al., (2018) compared to AF3. Furthermore, we have also compared the predictions from ESMFold and RoseTTAFold, where AF2 retained the lowest r.m.s.d. to resolved TREM2 structures. Then, we subsequently classified whether all missense mutations occur on high-confidence regions of the predicted structure. Using the predicted local distance difference test (pLDDT) score set above 70, we found 32 variants that are located on high-confidence residues of the 51 variants reported within prior literature. The results from this analysis have been provided in the **Supplementary Data**. Of the variants not included within our analyses, these residues occur on the cytoplasmic domain and stalk of the receptor. As a validation of the predictions obtained from AF2, we compared it to the partially resolved structure from Sodom et al., (2018) in the key IgV-like domain (PDB: 5UD7; Chain A).

### 2.2. Computational Tools

This study followed a similar protocol regarding the usage of computational tools to our recently reported work (Pillai et al., 2025). We exclusively used publicly-available softwares and web servers that have been peer-reviewed and validated by external studies. The workflow for this study has been illustrated in **Fig 1**. For predicting protein stability effects on a single protein chain, we relied on PremPS, DDMUT, DynaMut2, MAESTRO, DeepDDG, and INPS-MD (Chen et al., 2020; Zhou et al., 2023; Rodrigues et al., 2020; Laimer et al., 2015; Cao et al., 2019; Savojardo et al., 2016). Each algorithm differs from one another in approach, technique, and context for evaluating a sequence or structure of a protein, so we performed analysis of each result both individually and cumulatively. Next, we determined the functional effects of the mutations with PolyPhen-2, AlphaMissense, SIFT, and CADD (GRCh38v1.7) (Adzhubei et al., 2013; Cheng et al., 2023; Ng et al., 2003; Rentzsch et al., 2019). Structural analysis of the variants was demonstrated with Missense3D, as it remains the most comprehensive tool for reporting alterations in the protein structure. Lastly, all molecular visualization was completed with PyMOL (Schrödinger LLC, Portland, OR 97204, USA). We have provided the source files used for visualization described in the GitHub repository provided in the **Supplementary Data**.

**Figure 1.**
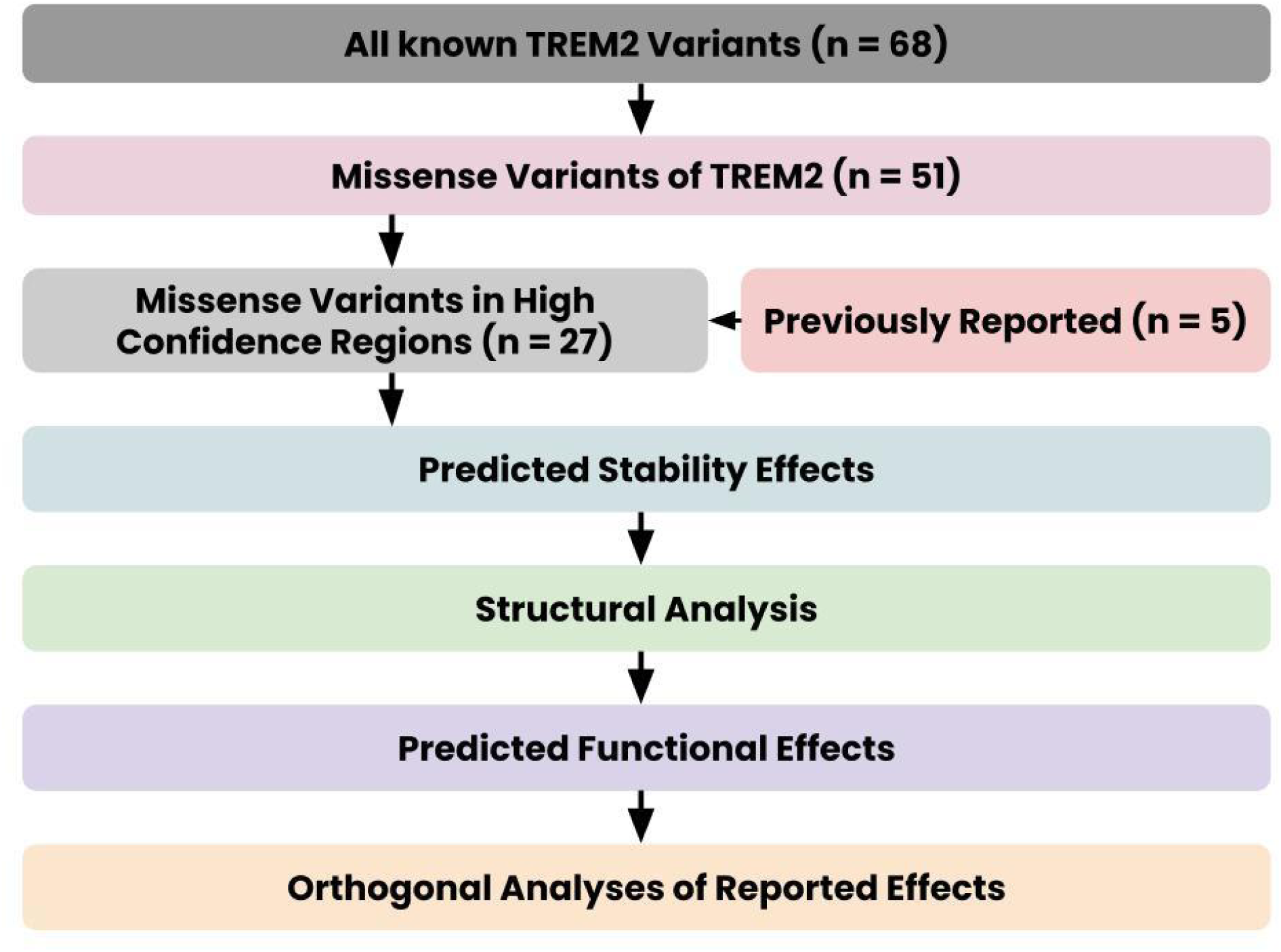
Workflow of this study for evaluating TREM2 variants. A simplified workflow for evaluating the TREM2 variants. There are 68 unique mutations that have been reported within known literature, provided on Alzforum.org. This count was reduced to 51 missense mutations, and subsequently to 32 because of adequate pLDDT score on the AF2 model of TREM2. Firstly, stability effects of each variant were predicted using the numerous computational tools. Then, structural analyses were performed of each variant using Missense3D. Lastly, functional effects were determined and orthogonal analyses were performed of the bioinformatic data.

### 2.3. Statistical Analyses

After collecting the source data from the in-silico algorithms, it was formally placed in a Microsoft Word document and subsequently processed in Excel. For visualization and analysis, GraphPad Prism (version 5.4) was used (GraphPad Software, La Jolla, CA, USA). All data described has been represented as the mean ± standard error of the mean (SEM). Unless stated otherwise, α = 0.05 and *p*-values are indicated in the corresponding figures and tables. ns: non significance; \**p* < 0.05; \*\**p* < 0.01; \*\*\**p* < 0.001; \*\*\*\**p* < 0.0001. For the stability analysis, we used a one-sample Student’s *t*-test (H_0_ = 0 kcal/mol) to determine significance. We assumed that the change in stability from in-silico mutagenesis for a specific variant would be derived from a normal Gaussian distribution. Shapiro-Wilks normality test confirmed that all mutations were normal except for the A28V and H43Y variants, so there is not enough evidence to reject the null that the data is derived from a normal distribution. For comparisons involving two groups, a Welch’s two-sample Student’s *t*-test was performed.

## 3. Results

### 3.1. Significant structural damages occur in NHD-causing TREM2 variants

There are 4 missense mutations (Y38C, W50C, T66M, and V126G) reported in literature that are likely NHD-causing variants (Kober et al., 2016). In our previous report, we provided predictions for the Y38C and V126G that were found to impose greater instability on the 3D structure compared to the AD-causing variants (Pillai et al., 2025). Herein, we predicted the molecular basis of the W50C and T66M mutations for completion. From structural analysis of the W50C variant with Missense3D, no significant damages were detected to the structure. The relative solvent accessibility (RSA) increased from 0% to 5.9% while the clash scores decreased from 17.96 to 13.73 (**Fig 3**). The WT residue was neither involved in hydrogen bonding nor the mutant residue. The structure of the WT and W50C variants have been visualized in **Fig 1A-B**. From stability analysis, the W50C variant significantly destabilized the structure by -1.583 ± 0.254 kcal/mol (*p* = 0.0016) (**Fig 1E**). According to all functional predictions, the mutant would be disruptive according to all 4 algorithms (**Table 1**). From the orthogonal data, it may be possible that despite decreased clash scores, the reduced bonding could destabilize the protein, eventually causing it to become non-functional. This was originally hypothesized by Kober et al., (2016) that the buried mutations could be negatively impacting protein folding of TREM2.

**Table 1.**
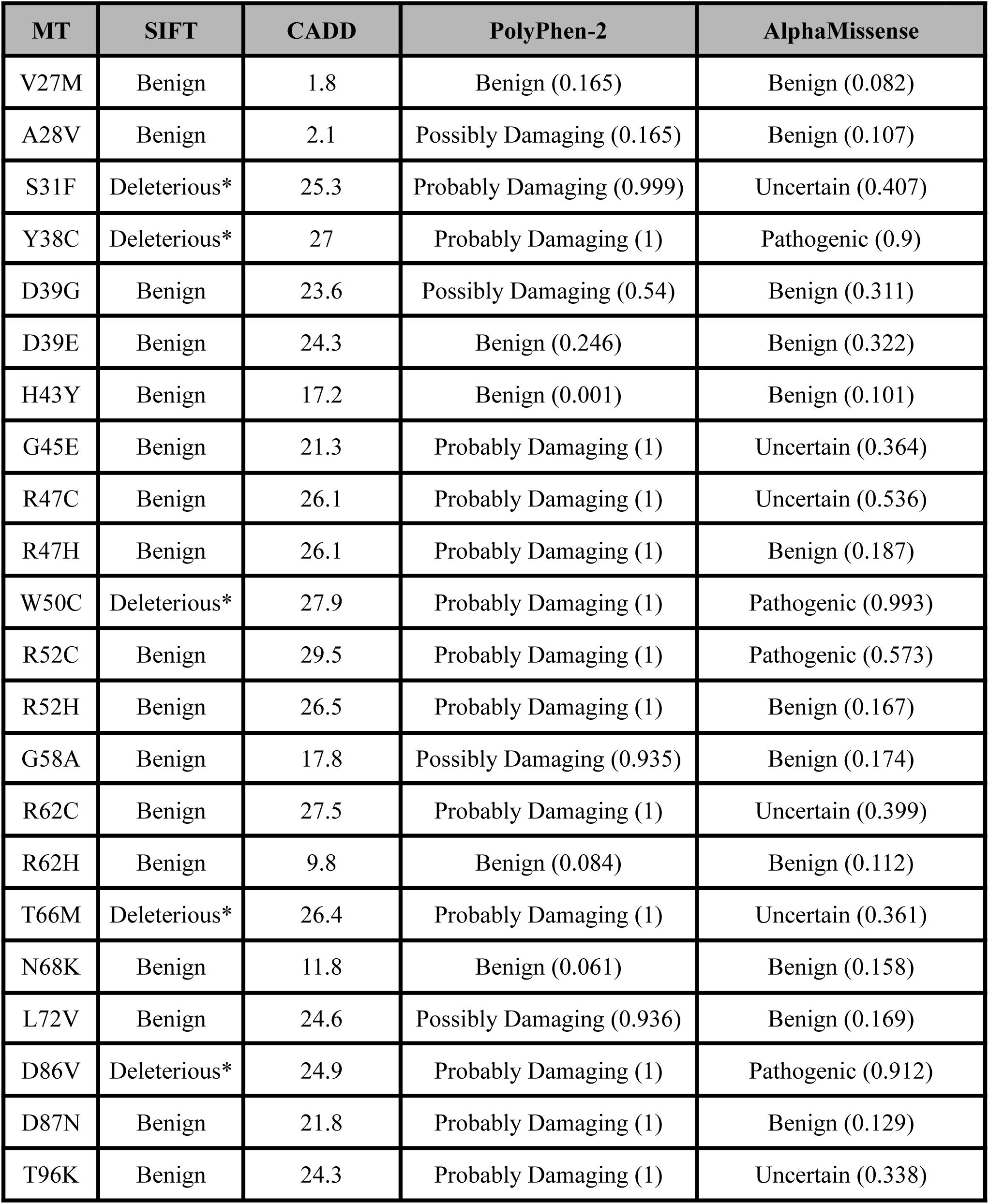
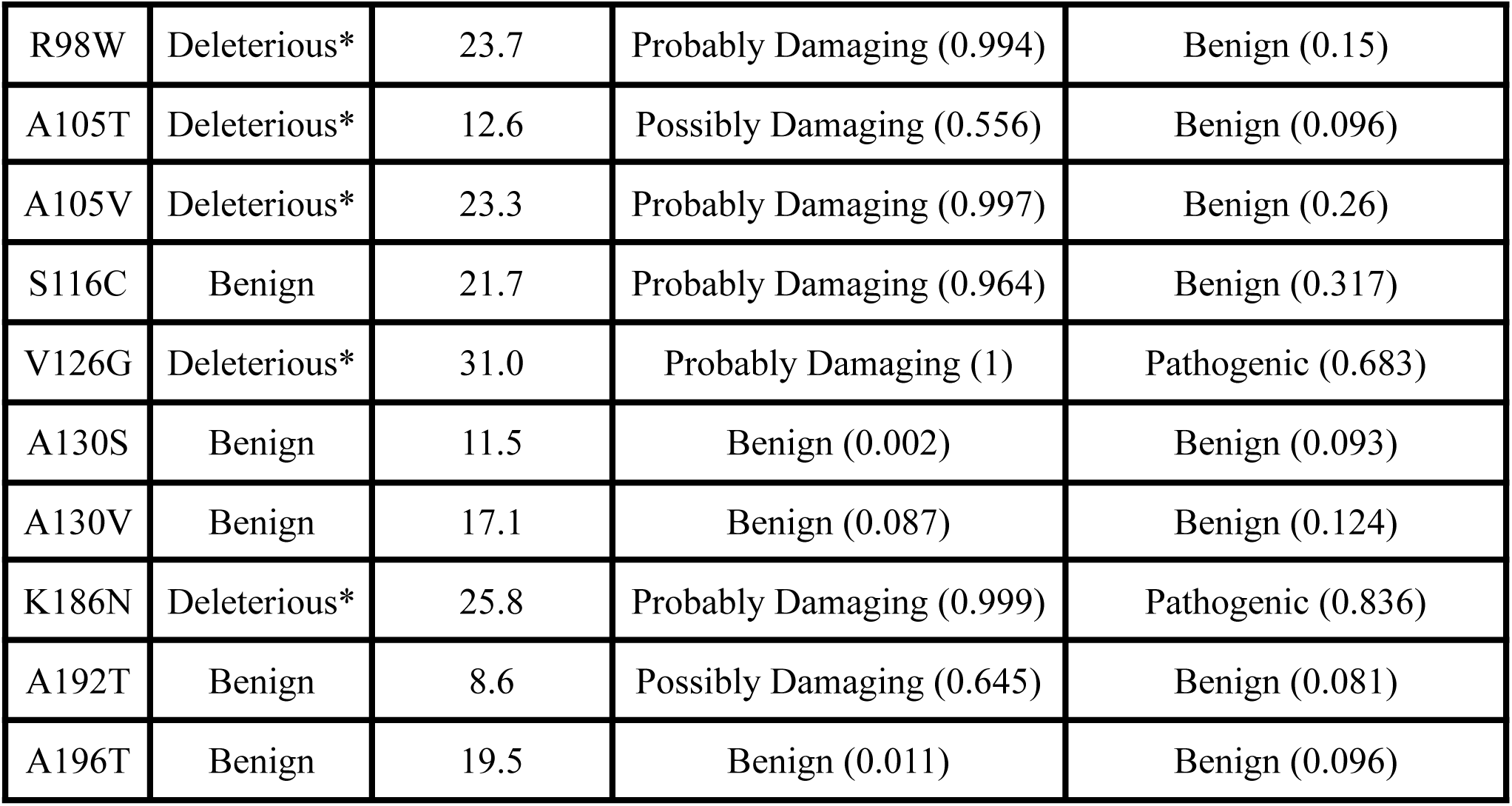
Pathogenicity of TREM2 variants using orthogonal bioinformatic approaches. AlphaMissense predictions were obtained on January 30, 2025 from the AlphaFold Structure Database. CADD predictions were taken on ProtVar (https://www.ebi.ac.uk/ProtVar/) using the UniProt accession identification (Q9NZC2) with the missense mutation. The genetic position of each variant was derived from Alzforum.org. CADD scores greater than 25 are considered deleterious variants, 20-25 are moderate, and below are benign. PolyPhen-2 predictions were derived from UniProt. *Low confidence in prediction by SIFT.

Next, we performed the same analysis with the T66M variant. From structural analysis, Missense3D had similarly predicted that no significant structural damages would occur on TREM2. Interestingly, the RSA had decreased from 0.7% to 0.5% and the clash scores had increased from 23.35 to 37.27 (**Fig 3**). In the WT structure, the carbonyl of Thr66 is involved in a hydrogen bond network with the nitrogen and amine of Arg47 and the hydroxyl of Ser65. There is also involvement between the hydroxyl of Thr66 and carbonyl of His67, carbonyl and nitrogen of Lys48 (**Fig 1C**). However, after site-directed mutagenesis, most of the hydrogen bond network is broken, with the carbonyl of Met66 interacting with the hydroxyl of Ser65 only (**Fig 1D**). Stability analysis determined that the variant causes significant destabilization to the structure -0.897 ± 0.235 kcal/mol (*p* = 0.0002). From a comparison of the W50C and T66M variants, it was found that the W50C variant imposed greater destabilization compared to the T66M mutation (*p* = 0.0426), which is surprising as T66M is involved in more critical hydrogen bonding and buried further in the structure. We hypothesize that based on the biochemical data, it appears that W50C variant causes greater solvent exposure compared to T66M, and could be a potential factor for greater destabilization to the structure of TREM2. From functional predictions of T66M, all algorithms expected the variant to be pathogenic except for AlphaMissense that was uncertain (**Table 1**).

In the context of the Y38C variant, the difference in destabilization of TREM2 was not significantly different to W50C (*p* = 0.8399), but was significantly more than that of the T66M variant (*p* = 0.0216). Of all variants of the receptor, the V126G variant caused the greatest destabilizing effect on the structure compared to all other variants (**Fig 1E**). Overall, for NHD-causing variants in TREM2, all appear to be buried in the structure and have critical stereochemical interactions occurring, which likely impact protein folding. In **Fig S1**, we received similar predicted results with the WT structure partially resolved by Sodom et al., (2018) for the W50C variants, and mostly similar to the T66M. On the WT experimental structure, interactions with Arg47, Lys48, and Ser65 are observed, but one additional interaction with Arg77 is noted. After mutagenesis, both interactions with Lys48 are missing but the remaining interactions are unchanged. The molecular details for the T66M variant still remain unclear given the differences observed between the two structures. It may also be possible that the T66M variant has less stability alterations compared to W50C because it still retains some hydrogen bonding after mutagenesis compared to the WT.

### 3.2. Evaluation of TREM2 variants in the transmembrane domain

Within this study, there were 3 variants that were located on high-confidence residues of the transmembrane domain, including the K186N, A192T, and A196T. From a structural analysis, all variants were predicted to not cause significant damage to the structure of the receptor and are located on the surface of the protein (RSA > 25%) (**Fig 3B**). The K186N variant was not involved in any hydrogen bond network, and no other relevant stereochemical effects were reported in its structure. However, the A192T variant did not have any interactions in its WT structure, but an interaction is observed between the hydroxyl of Thr192 and carbonyl of Leu188. Unfortunately, given that the transmembrane domain has not been resolved, no comparison exists of predictions of the A192T variant. This is a similar case to the A196T variant that had no interaction occurring in the WT structure, but an interaction occurs between the hydroxyl of Thr196 and the carbonyl of Ala192. Overall, the molecular structure for all 3 variants have been provided in **Fig S2**. From stability analysis, the K186N, A192T, and A196T all destabilized TREM2 by -0.115 ± 0.214 kcal/mol (*p* = 0.055), -0.224 ± 0.212 kcal/mol (*p* = 0.183), -0.268 ± 0.456 kcal/mol (*p* = 0.292) that were not significant from no change in stability, respectively (**Fig 2E**). Lastly, from functional prediction algorithms, all methods predicted that the K186N variant would be pathogenic while the A196T to be benign, and mixed results for A192T (**Table 1**).

**Figure 2.**
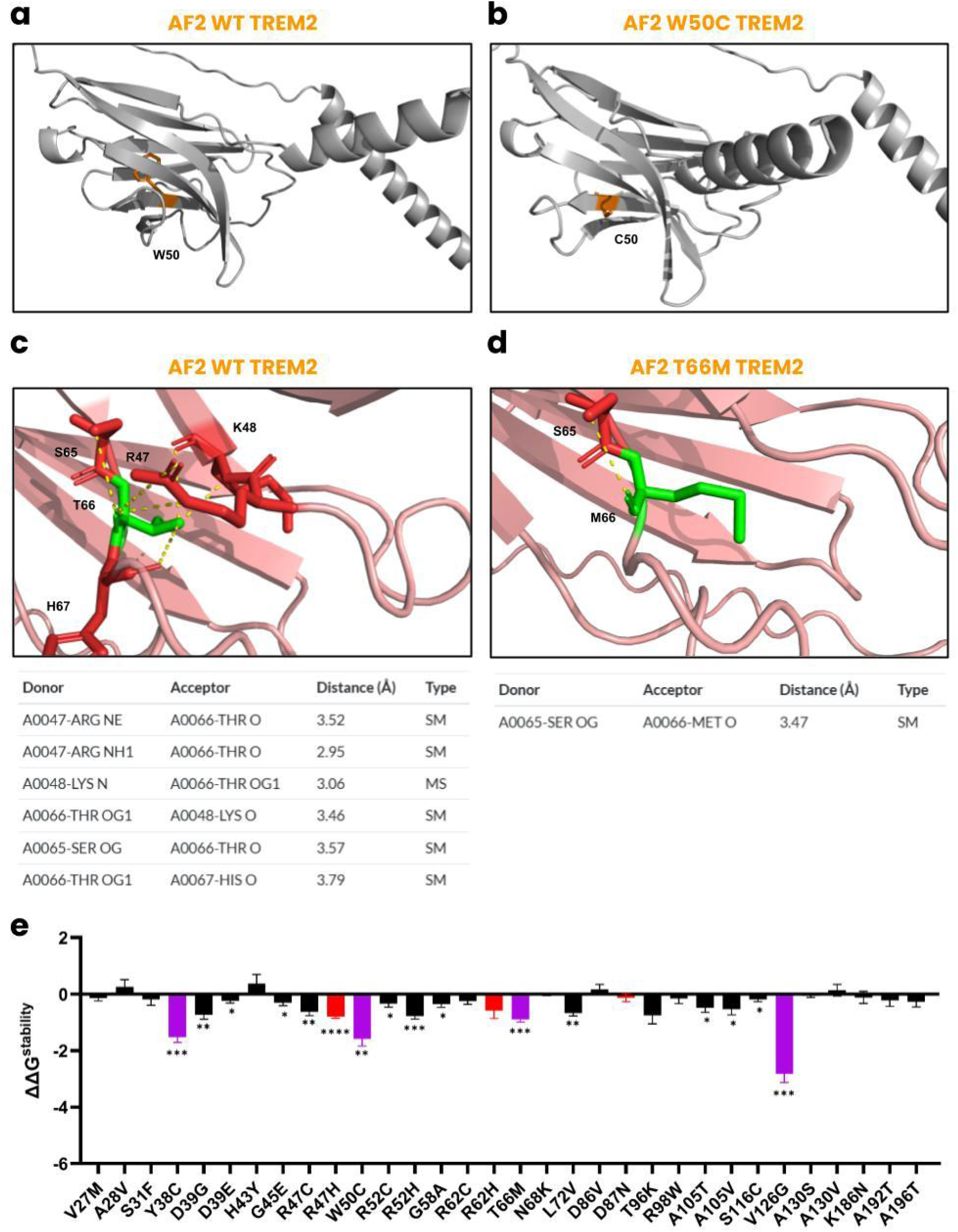
Predicted molecular basis of W50C and T66M TREM2 variants involved in NHD. **(a)** The TREM2 structure predicted by AF2 focused on Trp50 that is buried deeply in the structure and not involved in any hydrogen bonding networks with other residues. **(b)** After in-silico mutagenesis, the structure is focused on Cys50, where the mutant still retains structural similarity to the WT. **(c)** At Thr66 on the structure, there are hydrogen bond networks involved with Arg47, Lys48, and Ser65. **(d)** After mutagenesis, the structure is predicted to only retain an interaction with Ser65.

**Figure 3.**
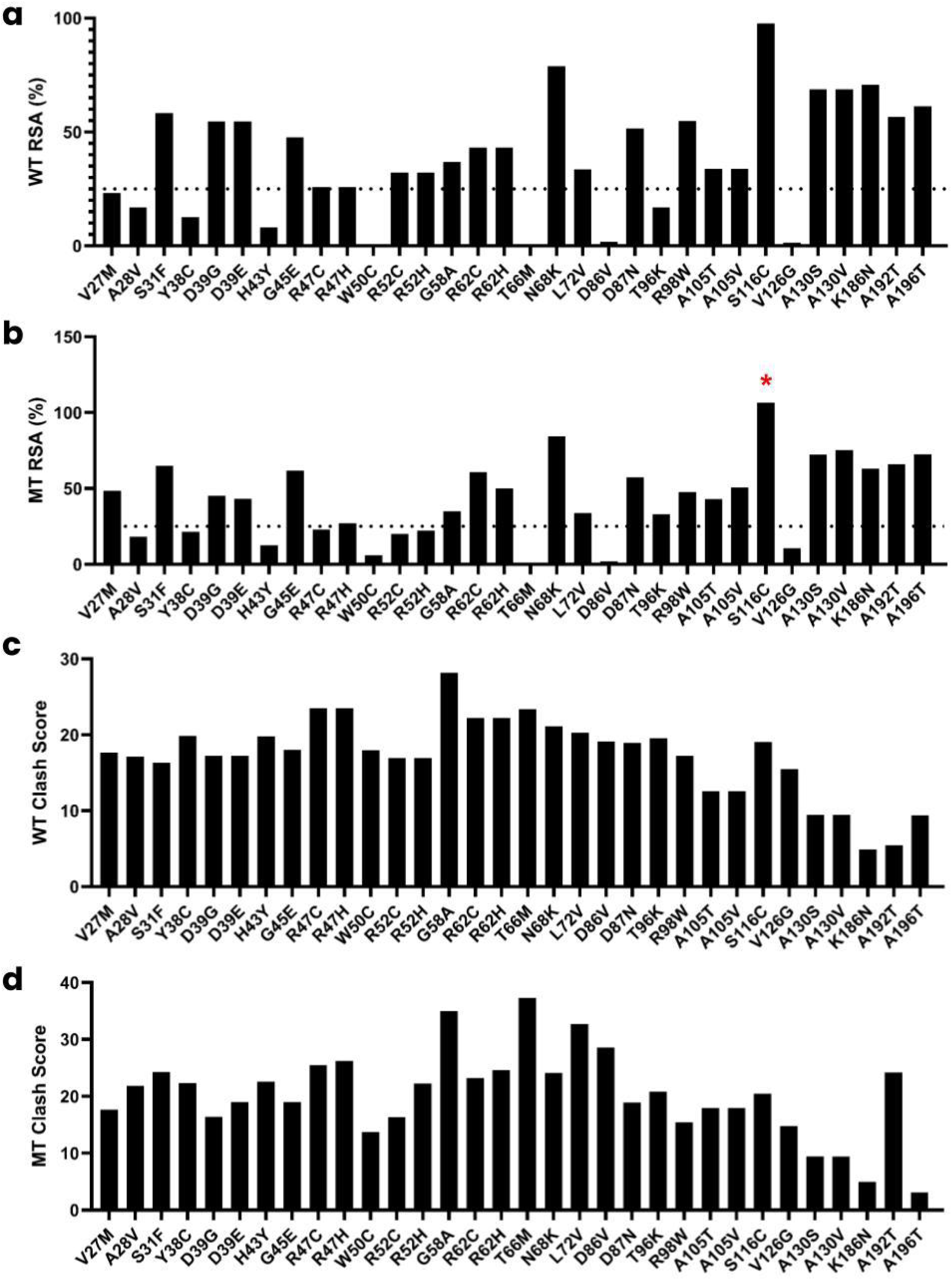
Structural analysis of all 32 TREM2 variants. **(a)** The percent of relative solvent accessibility (RSA) for the residues on the WT protein is provided in the bar graph. (b) Likewise, the RSA for the mutant residues is provided. **(c)** Clash scores for the WT residues are provided, and **(d)** those of the mutant are displayed. *The percent for RSA of the S116C variant had a score above 100% provided by Missense3D.

### 3.3. Predicted molecular details of TREM2 variants on the surface of IgV-like domain

The IgV-like domain is the most high-confidence region of the AF2 structure, ranging from residue 19 to 131. Given the large number of variants on this critical domain, we report variants located on the surface of the protein within this section. From the RSA above 25% on the WT structure, our analysis included the S31F, D39E, D39G, G45E, R47C, R47H, R52C, R52H, G58A, R62C, R62H, N68K, L72V, R98W, A105V, A105T, S116C, A130S, and A130V.

The R47H and R62H variants were not included in our analysis because their predicted molecular details have been reported previously (Pillai et al., 2025). Given the large number of variants on the surface of this domain, we have divided our analyses into residues 31-47, 52-68, and 105-130.

For the S31F variant, structural analysis revealed that no significant damages were made to the structure given that the residue did not have any hydrogen bond networks or other interactions involved, but there was increased clash scores and higher solvent exposure. Functional predictions provided mixed results for this variant. Lastly, from stability analysis, the receptor destabilizes by -0.185 ± 0.209 kcal/mol that was not significantly different from no change (*p* = 0.418). The molecular details of S31F are visualized in **Fig S3**. Next, for the D39E and D39G variants, both mutations resulted in lower RSA and local clash scores. D39E and D39G both significantly decreased stability by -0.242 ± 0.069 kcal/mol (*p* = 0.017) and -0.734 ± 0.155 kcal/mol (*p* = 0.0052). In the WT structure, Oδ1 Asp39 forms hydrogen bonds with the nitrogens of Met41 and Lys41, but loses both interaction after mutagenesis to Gly39, and only retains interaction with Met41 with Glu39 (**Fig 4**). Interestingly, most functional predictions expected both variants to be benign (**Table 1**). Similarly to the results of the S31F variant, we found that G45E increases clash scores and solvent exposure, but is not involved in any hydrogen networks involving other residues (**Fig S3**). Yet, the mutation did significantly destabilize the structure by -0.303 ± 0.105 kcal/mol (*p* = 0.034), and mixed results were obtained from functional predictions. To conclude, we examine the R47C variant and how it differs from the influential R47H mutation. As previously reported (Sodom et al., 2018; Pillai et al., 2025), Arg47 interacts with Ser65, Thr66, His67, and Asn68, but after in-silico mutagenesis Cys47 only interacts with His67 (**Fig 4**). R47C significantly destabilizes the protein by -0.628 ± 0.130 kcal/mol (*p* = 0.0048), and lesser but not significant from that of R47H (*p* = 0.292). Functional predictions are nearly similar for both variants. Given the biochemical data for the R47H and R47C, the molecular mechanisms remain unclear for how R47C leads to FTD as opposed to R47H that causes AD, and will be discussed in context of current experimental studies in **Section 4** Overall, we have also provided validations of the 5 variants using in-silico mutagenesis on the experimental structure by Sodom et al., 2018 in **Fig S4**. All experimental structures matched the interactions aforementioned except for the mutant single interaction with Met41 on D39E.

**Figure 4.**
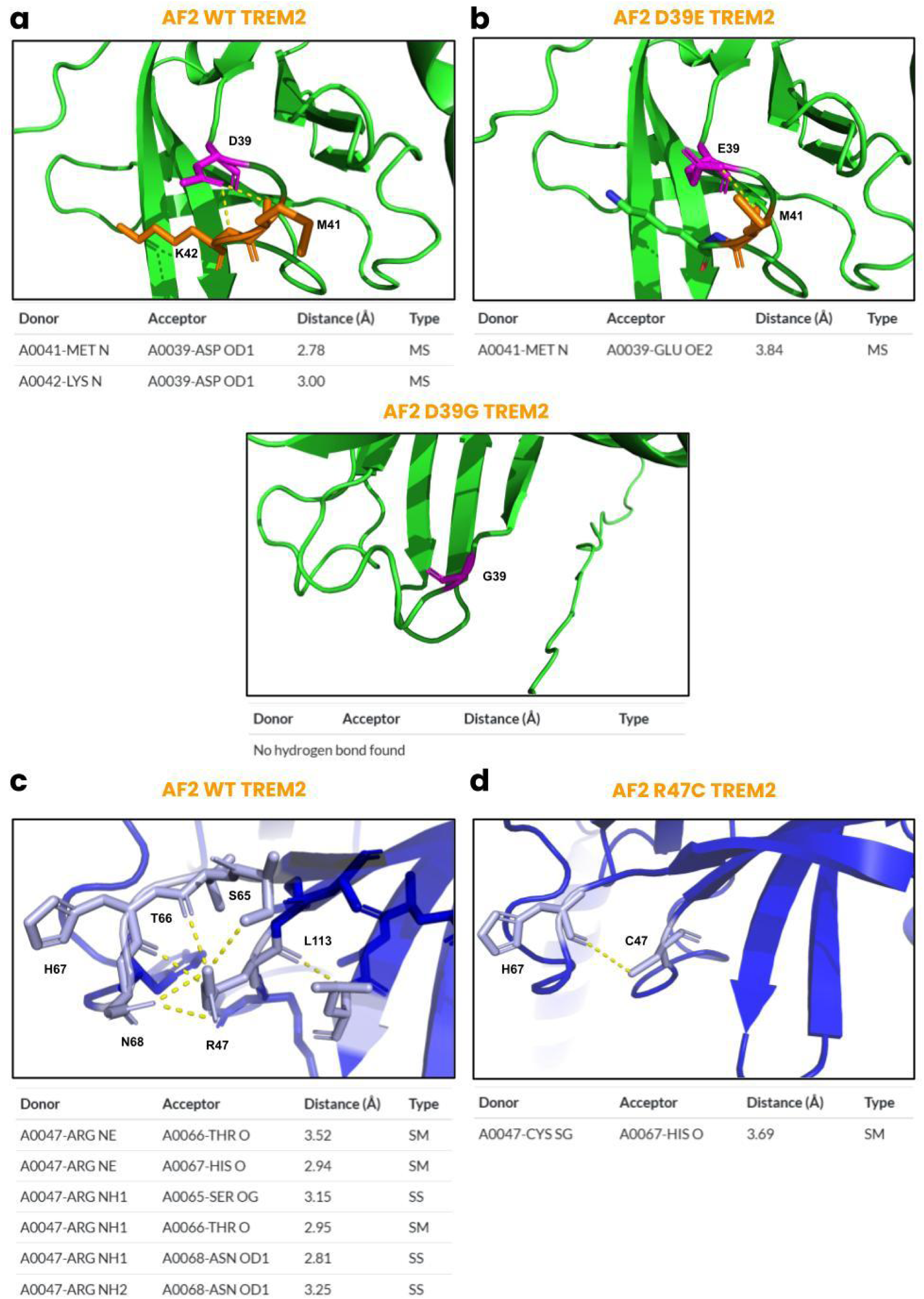
Predicted molecular details of D39E, D39G, and R47C. **(a)** Asp39 is expected to interact with the nitrogen of Met41 and those of Lys42, and **(b)** after in-silico mutagenesis the interaction with Met41 is retained in D39E and none in D39G, respectively. **(c)** Arg47 interacts with Ser65, Thr66, His67, and Asn68, which has been previously described in Pillai et al., 2025. (d) After mutagenesis, Cys47 only retains its interaction with His67.

Next, we evaluated the R52C and R52H variants. Both mutations cause significant destabilization of the TREM2 structure, with -0.338 ± 0.122 kcal/mol (*p* = 0.0391) and -0.776 ± 0.108 kcal/mol (*p* = 0.0008) for R52C and R52H, respectively. Although both variants do not cause significant structural damages to the receptor, clash scores and RSA remained relatively similar levels. Functional predictions provided mixed results, but most expected the variants to be pathogenic (**Table 1**). In the WT structure of Arg52, the two amine groups of the residue interacts with the carboxyl group of His103 and Ala105 (**Fig 5A**). After in-silico mutagenesis, both His52 and Cys52 are expected to not be involved in any hydrogen bond interactions. Our analysis then continued to the G58A variant, where we found significant structural damages from the mutation, where the residue was originally located in a bend curvature, and its replacement substantially increased local clash scores. This variant also caused significant destabilization to the receptor, but was significantly smaller at -0.344 ± 0.124 kcal/mol (*p* = 0.039), and is not involved in any hydrogen bond interactions and functionally expected to be benign (**Fig 5C**; **Table 1**). Following the G58A variant, we determined that the R62C variant did not significantly destabilize TREM2 by -0.247 ± 0.113 kcal/mol (*p* = 0.0817). This variant had also broken the expected hydrogen bond between the amine of Arg62 with carbonyl oxygen on Gln111 that was predicted from our prior report (Pillai et al., 2025), but its functional effects were uncertain from the algorithms (**Fig 6**). We then evaluated the N68K variant, which appeared to impose almost no effects on the structure. From stability analysis, the mutation caused destabilization by -0.0029 ± 0.044 kcal/mol (*p* = 0.9501), and all functional prediction algorithms expected the variant to be benign. In the WT structure, Oδ1 Asn68 interacts with the amines of Arg47 that have been previously confirmed with R47C, and after mutagenesis is both removed (**Fig 6**). Our analyses then continued to L72V, where structural analyses revealed that non-significant damages were made to the receptor as the residue is not involved in any other hydrogen bond networks (**Fig 6E-F**). Likewise, most algorithms expected the variant to be benign but destabilization was significant at -0.672 ± 0.102 kcal/mol (*p* = 0.0012). Overall, we have performed in-silico mutagenesis of these variants, and found that all residues involved in interactions were similar except for the mutant structure of R52C, where C52 interacts with Leu107 (**Fig S5**).

**Figure 5.**
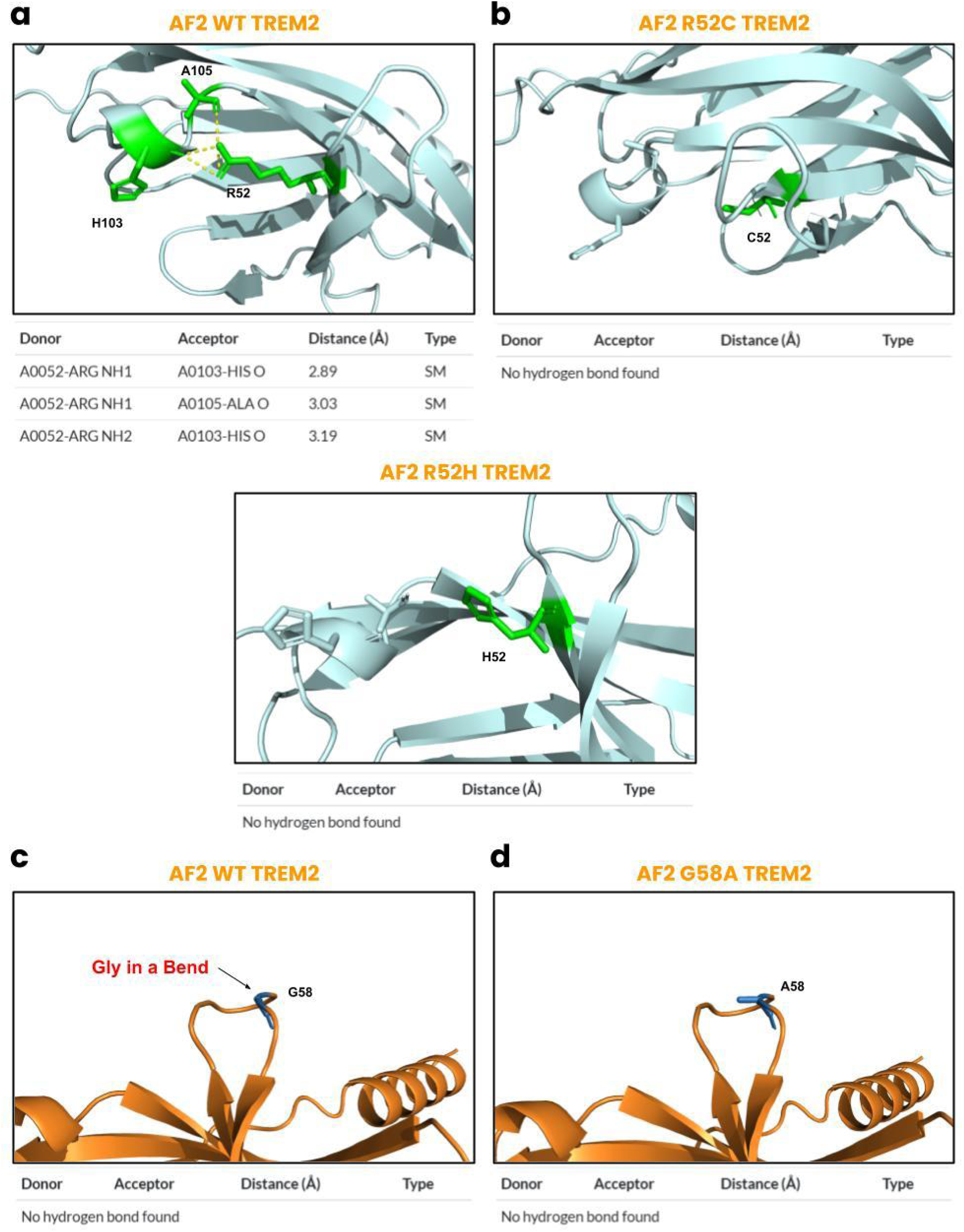
Predicted molecular basis of R52H, R52C, and G58A. **(a)** The amine of Arg52 forms interactions with the oxygen atoms of His103 and Ala105, **(b)** but was missing in both the replacement of R52C and R52H variants. **(c)** Gly58 is not involved in any interactions of other residues, **(d)** but is in a bend that is disrupted after mutagenesis of the AF2 structure.

**Figure 6.**
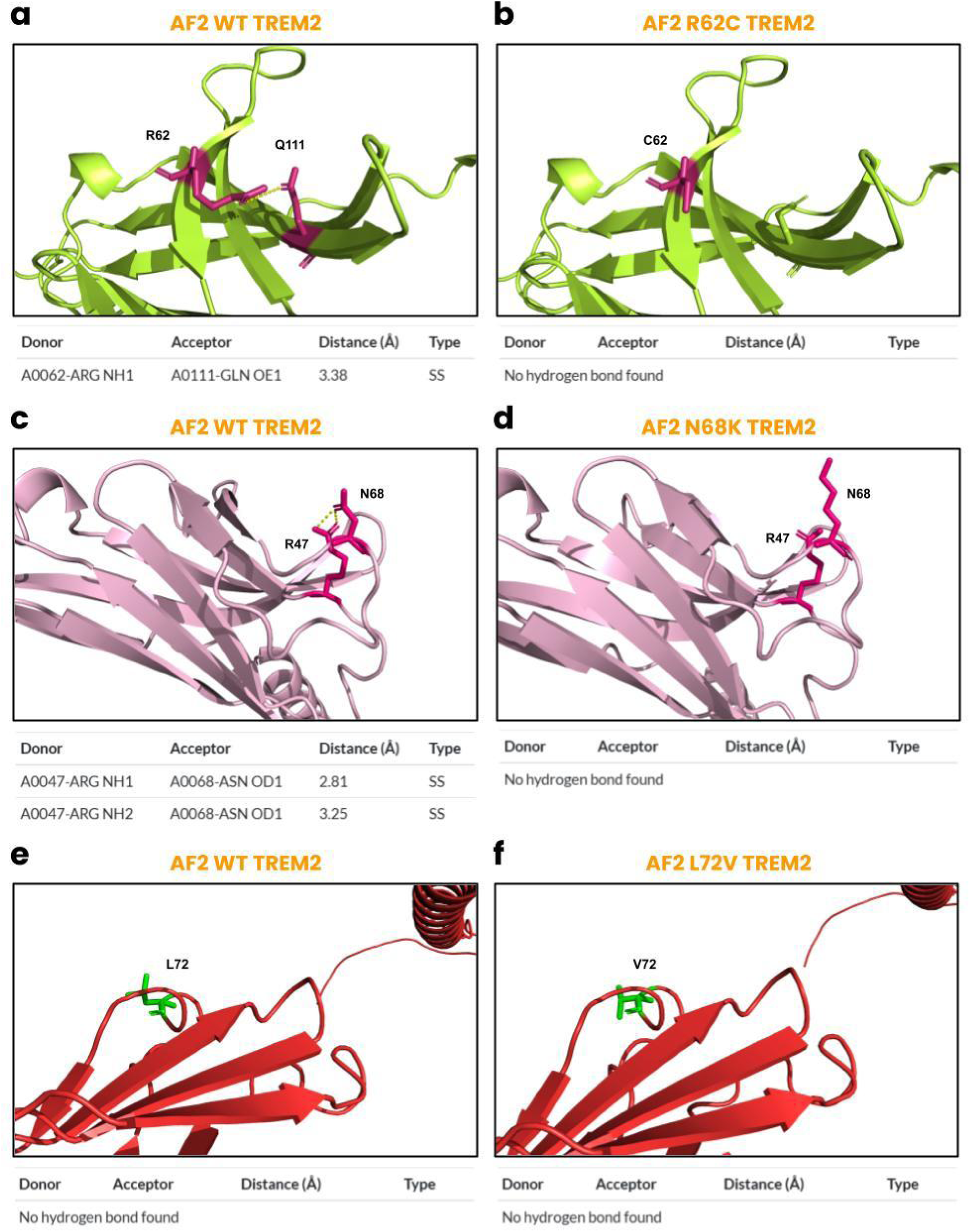
Molecular structures of R62C, N68K, and L72V. **(a)** Arg62 is involved in an interaction with Gln111, **(b)** but this interaction is removed after in-silico mutagenesis. **(c)** Similarly, Asn68 is involved in two interactions with Arg47, **(d)** but is removed in the predicted mutant structure. **(e-f)** Conversely, the L72V variant was not involved with other residues on the surface of the receptor.

In our final cluster of analysis, we studied the R98W, A105V, A105T, S116C, A130S, and A130V variants of TREM2. The R98W variant revealed that the variant does not disrupt the hydrogen bond interaction with Ser81 (**Fig 7A-B**), and the functional results were mixed and stability change was not significant at -0.157 ± 0.176 (*p* = 0.413). In the WT structure, Ala105 only interacts with amine of Arg52. However, the A105V variant results in its removal and the A105T variant results in two hydrogen bonds between the hydroxyl of Thr105 and Pro102 and Val128 (**Fig 7C-D**). Both variants also caused significant destabilization, where A105T was -0.480 ± 0.162 kcal/mol (*p* = 0.0315) and A105V was -0.534 ± 0.197 kcal/mol (*p* = 0.0425), respectively. Functional results were mixed for both mutations **(Table 1)**. Next, we evaluated the S116C variant, where no significant structural damages were observed and the residue was not involved in any interactions (**Fig S6**). Yet, the mutation did result in significant destabilization of -0.192 ± 0.074 kcal/mol (*p* = 0.0493), and most functional predictions were benign. We then analyzed A130S and A130V, where both variants did not involve any interactions and all functional predictions were unanimous that the mutations are benign. Furthermore, no significant changes occurred in protein stability for Ser130 by -0.063 ± 0.053 kcal/mol (*p* = 0.227) and Val130 by 0.143 ± 0.203 kcal/mol (*p* = 0.541), respectively. The molecular details of both variants have been visualized in **Fig S6**. After performing in-silico mutagenesis on the partially resolved experimental structure, the results matched for WT Arg98, S116C, A130V, and A130S, and WT Ala105. However, both variants of A105 structure had retained the interaction with Arg52, and A105T instead gained a hydrogen bond with Val126 instead of Val128. Furthermore, the hydrogen bond interaction between Ser81 and Tyr98 was missing in the mutant structure (**Fig S7**).

**Figure 7.**
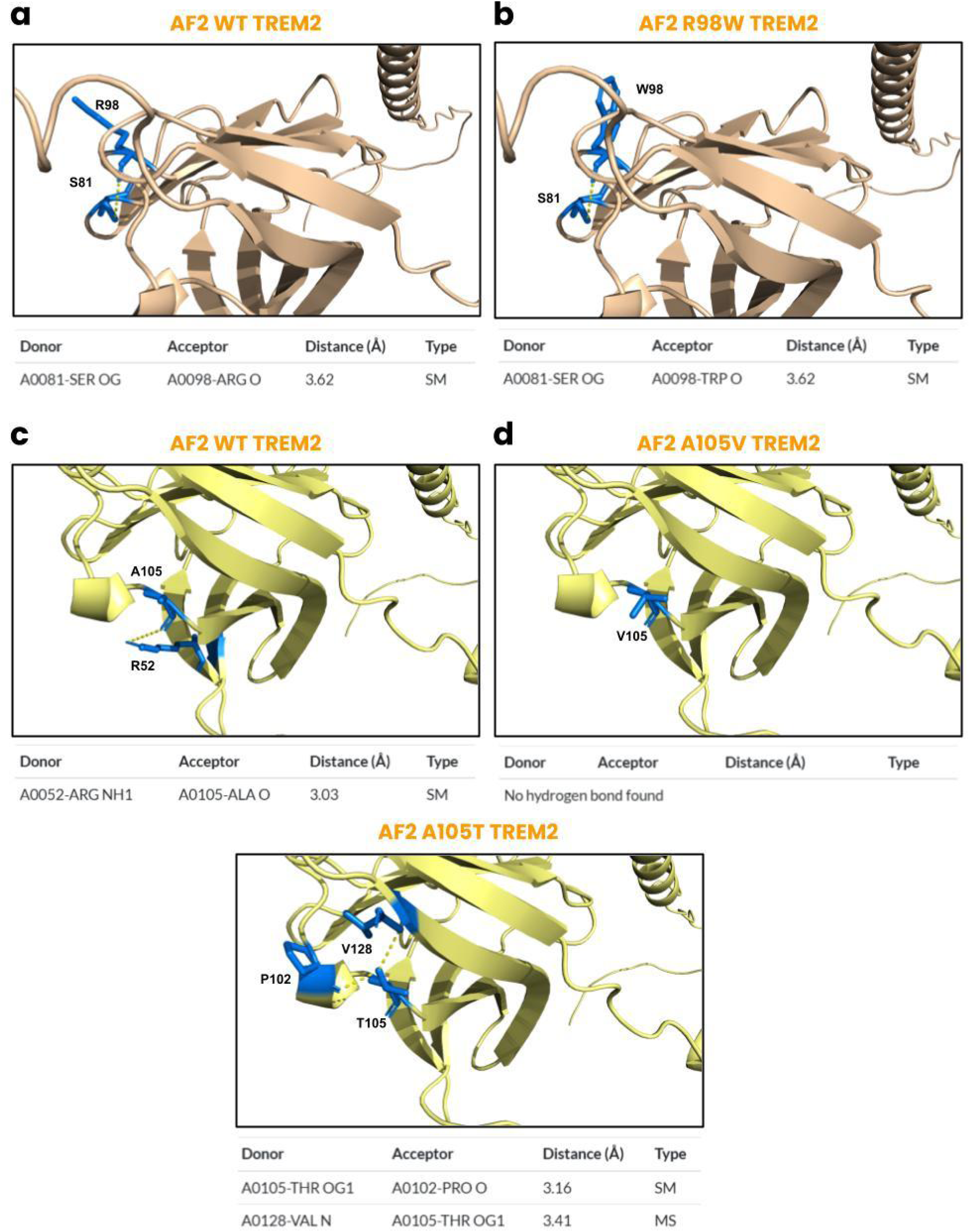
Predicted molecular structure of R98W, A105V, and A105T variants. **(a)** On the WT structure, Arg98 interacts with Ser81 that is **(b)** retained after replacement to Tyr98. **(c)** Similarly, on the structure of Ala105, interactions are observed with Arg52, **(d)** but none are retained in A105V and novel interactions are formed between Thr105 with Pro102 and Val128.

### 3.4. Predicted molecular details of buried variants in the IgV-like domain

Given the large number of variants on the IgV-like domain of TREM2, we split our analysis into those on the surface and buried in the structure. In this section, we evaluated residues located buried in the receptor (WT RSA < 25%), which included the V27M, A28V, H43Y, D86V, D87N, and T96K. We excluded T66M, V126G, Y38C, and W50C in our analysis as these variants have been previously evaluated.

Our first analysis included the V27M variant. From structural analysis, the variant did not impose significant damages to the structure, and was not involved in any notable hydrogen networks with other residues (**Fig 8A-B**). Similarly, Met27 did not significantly destabilize the receptor with -0.145 ± 0.096 kcal/mol (*p* = 0.192), and was unanimously classified as benign by all functional algorithms. Next, we evaluated A28V that yielded similar structural results to V27M. There are no expected notable interactions involved in both the WT and mutant structure (**Fig 8C-D**), and most functional predictions expected the variant to be benign. Val28 also did not result in significant destabilization of TREM2 at 0.259 ± 0.251 kcal/mol (*p* = 0.313). Interestingly, we observed a contrast to this trend with the H43Y variant. By replacing His43, the interactions between the nitrogen from the imidazole ring and the side chain hydroxyl of Ser112 is removed, and the salt bridge formed between the imidazole ring and the carboxyl group on Asp119 is disrupted (**Fig 8E-F**). Yet, all functional algorithms were unanimous that the variant is benign and the change in stability was not significant at 0.369 ± 0.329 kcal/mol (*p* = 0.436). We hypothesize that the replacement of H43Y improves the stability of the receptor noticeably but not significantly, and therefore was expected to be benign. We also observed another exception to the general trend with D86V, where significant damages were made to the receptor as it is replaced with an uncharged residue and all hydrogen bond interactions involved between the side-chain carboxyls of Asp86 and the hydroxyl of Tyr38 and terminal nitrogen on Lys42 (**Fig 9A-B**), and all functional predictions expected the Val86 to be damaging. Despite causing damage, its destabilizing effects were not significant at 0.165 ± 0.181 kcal/mol (*p* = 0.404). The T96K mutation also followed the general trend observed in most variants, where no significant structural damages were detected and the residue was not involved in any interactions (**Fig 9C-D**), but functional predictions were mixed and change in free energy was not significant at -0.754 ± 0.301 kcal/mol (*p* = 0.055). Finally, we performed the in-silico mutagenesis on these mutations, where we confirmed the V27M, A28V, mutant structure of H43Y and D86V, and the T96K variant. However, on the WT structure of H43Y, no salt bridge was detected and an additional interaction with Arg46 was observed, and similarly for D86V with an additional interaction of Asp87 **(Fig S8)**.

**Figure 8.**
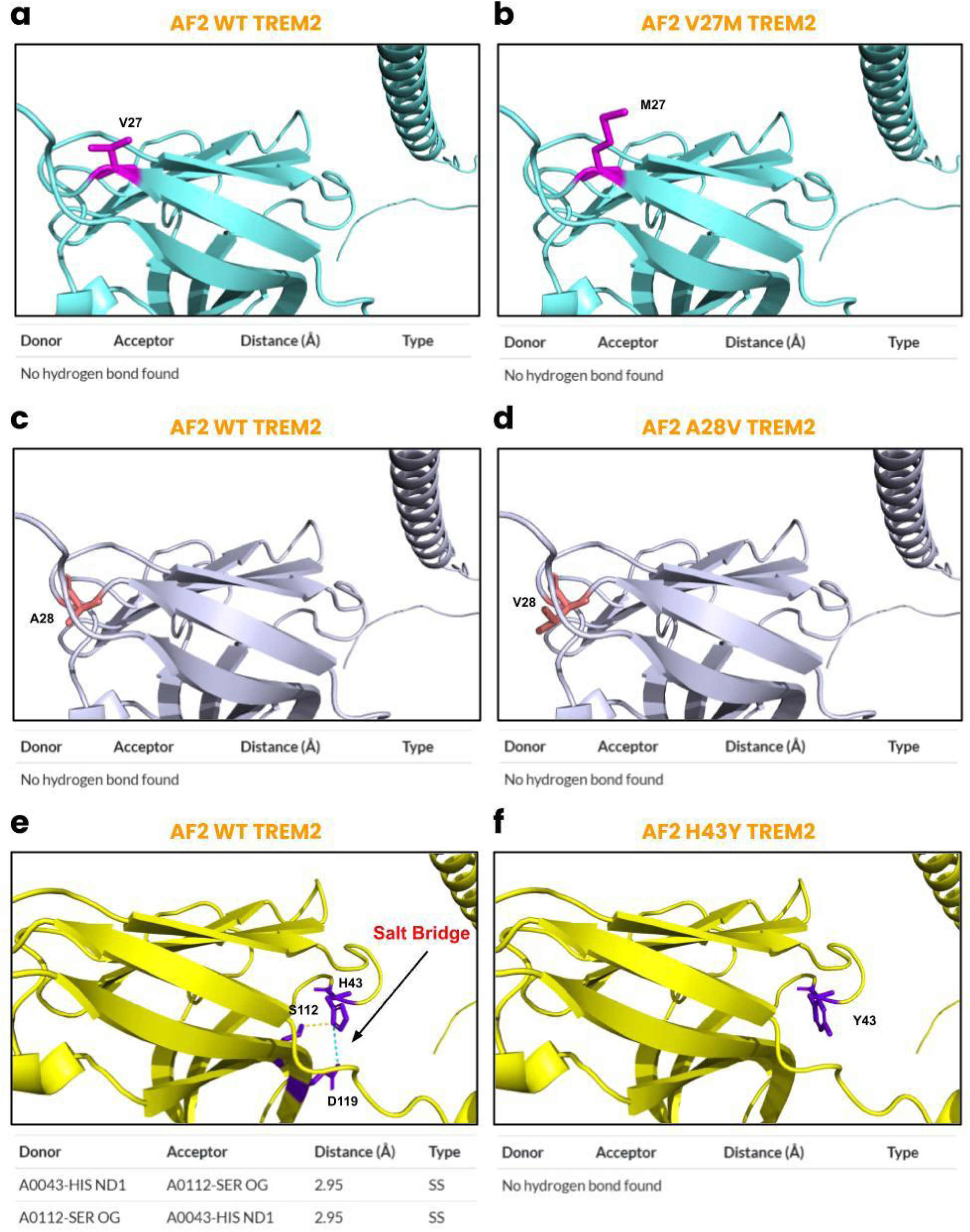
AF2-predicted molecular structures of V27M, A28V, and H43Y. (a-b) There are no interactions on the WT and mutant structures for the V27M variant. **(c-d)** A similar trend was observed with A28V, where no interactions were involved in either structures. **(e)** On the WT structure, His43 forms a hydrogen bond with Ser112 and has a salt bridge with Asp119 (visualized in cyan), **(f)** which are both removed in the mutant structure.

**Figure 9.**
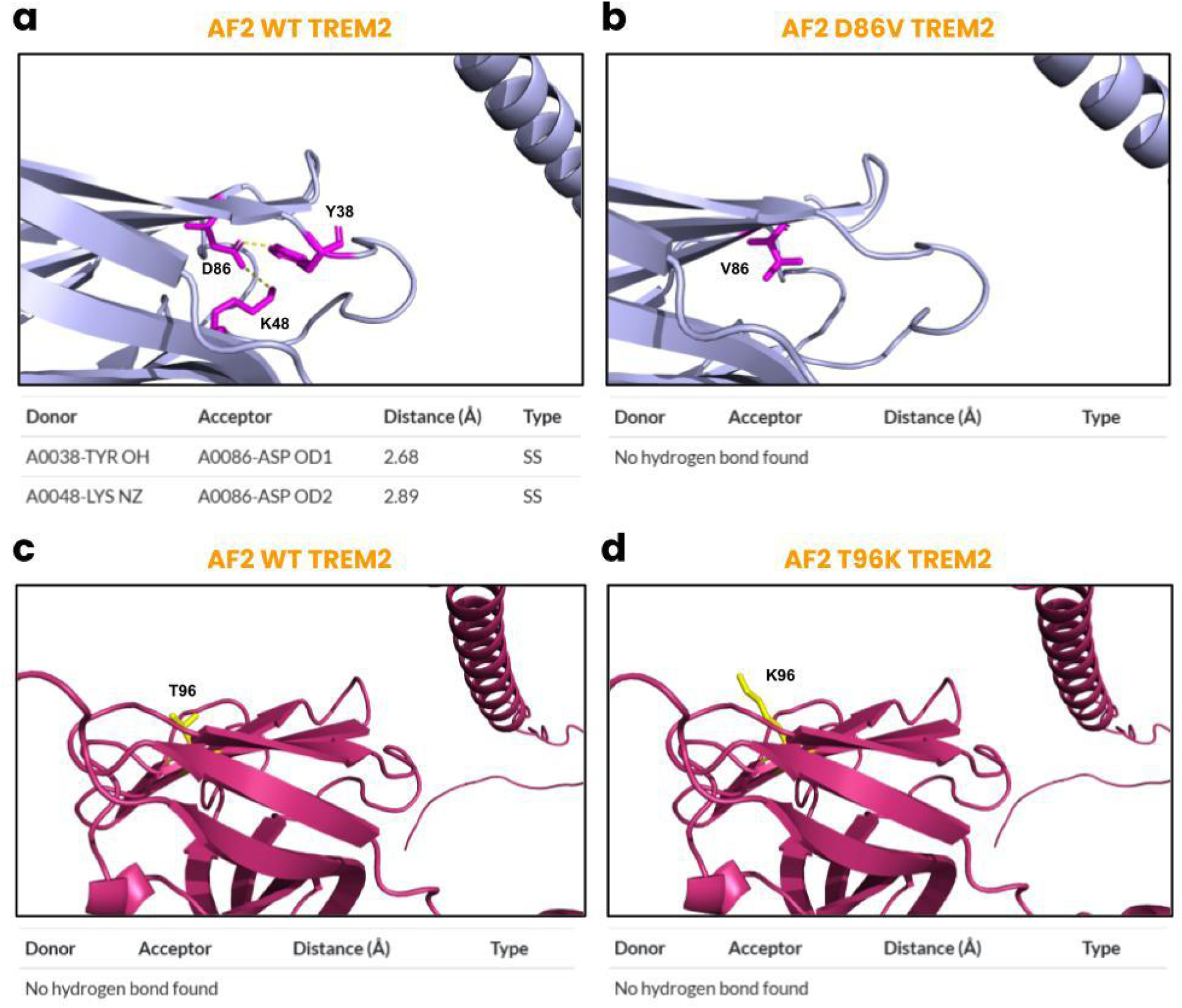
Predicted molecular details of the D86V and T96K. **(a)** In the WT structure, Asn86 interacts with Tyr38 and Lys48, **(b)** but after in-silico mutagenesis both interactions are removed. **(c-d)** Thr96 is neither involved in hydrogen bond interactions nor is the mutant structure.

## 4. Discussion

In the current study, we have expanded on our previous investigation (Pillai et al., 2025) and provided baseline biochemical data on most known missense variants of TREM2 reported in literature. We used our protocol to define the expected stereochemical, stability, and functional effects of variants involved in NHD and AD, and other miscellaneous variants located on the IgV-like domain and transmembrane domain of the receptor, cumulating in 32 variants. To our knowledge, this is the first study to report on the 3 variants involved in the transmembrane domain. Furthermore, we used our data to compare individual variants and to characterize their subtle differences, namely those implicated in NHD. We believe that the in-silico data provided in this study may be informative to researchers and clinicians alike towards improving understanding of TREM2 variants of unknown clinical and biological effects, and guiding experimental and clinical studies involving this crucial receptor in the future.

Within known literature, the four variants (Y38C, W50C, T66M, and V126G) implicated in NHD have been well-characterized. Interestingly, we found that while the W50C variant did lead to significant destabilization of TREM 2, there were reduced steric clashes in the structure. In comparison to W50C, T66M had a drastic increase in clash scores, but still caused less significant damage to the protein. These findings are directly supported by extensive molecular dynamic studies that were performed on the partially resolved structure from Kober et al., 2016 (PDB: 5ELI), where all variants except W50C were found to increase steric clashes and loss of ligand-binding affinity (Dash et al., 2020). With support of experimental studies from Kober et al., 2016, we put forth the hypothesis that NHD-causing variants cause the most damage to protein folding and stability of TREM2 in comparison to all other reported variants. Yet, it remains unclear of the mechanisms by which the W50C variant uniquely affects the receptor aside from the residue being buried. Of note, this variant is second to V126G in increasing the expected RSA of the residue, and could be a potential cause for its moderate destabilizing effect. Ultimately, to the best of our knowledge, this is the first study to provide orthogonal in-silico analyses of all NHD-causing variants, confirming prior experimental and simulation studies, as well as defining the W50C variant that remains unclear in current literature.

Herein, we also provide the predicted molecular details of variants on the transmembrane domain that have not been reported. Within literature, there is a poor understanding of the K186N, A192T, and A196T variants. In a prior study, the K186N variant was found to undergo normal cell maturation and high levels of cell expression, and was predicted to form defects to signal transduction with DAPI12, but not confirmed experimentally (Sirkis et al., 2017). In this study, the K186N variant did not impose significant alterations in stereochemistry and stability to the receptor. Likewise, the A192T mutation has been reported to reduce cell surface expression, but no other experimental studies have been completed (Bonham et al., 2017). Similarly to the results of the K186N variant, A192T did not significantly impact the receptor. Lastly, there has been no experimental studies in literature on any biological effects of the A196T variant other than its original reporting (Sirkis et al., 2016). From our predicted data, none of the WT structures for the transmembrane variants appear to be involved in hydrogen bonding interactions, but after in-silico mutagenesis, interactions appear with the A192T and A196T. The results of these variants are unique as the stalk and cytoplasmic domain are poorly defined in the 3D structure, due to the challenge to resolve experimentally through crystallography techniques. Overall, our current study provides baseline biochemical data for these variants of which no experimental efforts have been directed towards.

Now, the primary analyses involved in this study were variants located on the IgV-like domain, that either reside on its surface or are buried in the structure. For those on its surface, we notably highlight the D39G, D39E, R47C, R52C, R52H, G58A, L72V, A105T, A105V, and S116C variants for causing significant destabilization to the receptor. To our knowledge, there are no studies that have reported biological effects of the D39G and D39E variants. Given our in-silico data, we suggest that the effects of these two variants appear to be subtle as the residue is not involved in a complex network of interactions nor is it buried in the structure of the receptor. Next, we were interested in the effects of the R47C variant as this residue is critical for the influential R47H variant associated with AD. The data met expectations of this mutation because of the critical Arg47, where we found the variant destabilizes the structure, reduced interactions in the hydrogen bond network, and possibly damaging functional effects. However, it still remains unclear as to how the replacement of Cys47 does not lead to similar pathogenic effects as those of His47. We posit that Cys47 causes significant alterations to the 3D structure that differ from the rotations that were observed in His47 (Sodom et al., 2018). In a biophysical mapping study of TREM2, it was reported that effects on Arg47 as a basic site partially reduces interactions with ApoE and IL-34, but does not impact TDP-43 and C1q (Greven et al., 2025). From a recent pre-print from the same group, structural studies found that Aβ(1-8) binds with the hydrophobic sites of the CDR1 loop, but when mutations occur in the CDR2 loop, there is a reduction in binding of oligomerized Aβ42, and ultimately suggests that the conformationally dynamic CDR2 loop influences the ability of CDR1 to obtain a optimal conformation that directly binds to oAβ42 (Greven et al., 2024). With account of these previous structural studies and our in-silico biochemical data, it may be plausible that Cys47 does not severely affect the conformational dynamics between the CDR1 and CDR2 loops as opposed to His47. Further studies are required to explore the differences in molecular dynamics between the R47C and R47H variants.

Continuing to R52H and R52C, these variants like many others do not have biological or neuropathological effects reported thoroughly. In one experimental study, it was found that the R52H variant decreases cell surface expression compared to the WT in a reporter cell line, but this trend was not observed in those activated by phospholipids (Song et al., 2017). Even more limited was R52C that only had one clinical case study with a phenotype of xeroderma pigmentosum and behavioral patterns consistent with FTD (Soares et al., 2020) and no experimental studies. Yet, our findings for both variants is that significant hydrogen bond interactions are lost from both variants that lead to significant instability in the receptor, although functional results remain uncertain overall. Conversely, the G58A variant was one of the few structures that caused significant stereochemical effects to TREM2 as there was a bend occurring the curvature, and also resulted in destabilization of the structure. Although there are no experimental studies for this variant, a study identified this variant in 1216 AD patients and 0 FTD or 0 healthy Belgian individuals (Cuyvers et al., 2013). Next, the L72V variant was found to increase clash scores and cause destabilization, but was not involved in any interactions with other residues. We suggest that the L72V variant causes subtle but not profound effects on the receptor, as current clinical data has only identified this mutation in healthy individuals in a GWAS of a North American sample (Ghani et al., 2016). Similarly, with A105T and A105V, the mutations impose subtle effects on the structure although functional results remain uncertain. The A105T variant has only been reported in one study evaluating mutations in FTD subtypes (Thelen et al., 2014), while A105V was only confirmed to not be associated with AD in two clinical studies (Sims et al., 2017; Jin et al., 2015). Lastly, we evaluated the S116C variant on the surface of the domain, where results were similar in trend to the L72V, as there is subtle destabilization but no noticeable damage or functional effects hindered from the replacement. The similarity may also be relevant with clinical data, where it was found in only 1 patient with FTD and none in healthy and AD patients (Borroni et al., 2013), but experimental data suggests that cell surface expression was similar to that of WT TREM2 (Varnum et al., 2017).

We further analyzed TREM2 variants that were buried residues in the IgV-like domain. Interestingly, our data suggests that most of these mutations do not impose major structural alterations. With V27M and A28V, we denote that the stability and structural effects were not significant and functional predictions were unanimous that V27M was benign and A28V was mixed, respectively. The V27M variant has only been found in 1 AD patient and exhibited normal protein maturation, and A28V was found in 1 FTD patient and normal cell maturation and increased cell-surface expression (Sirkis et al., 2016). A similar trend was observed with the T96K variant in results, and supported by prior literature that the variant does not alter protein folding or protein aggregation (Kober et al., 2016), and lowers cell-surface expression compared to the WT (Song et al., 2017). We also reported on the H43Y and D86V variants that imposed significant structural alterations that differ from the general trends observed from prior point mutations discussed. H43Y was unanimously expected to be benign, but structural damages were detected as a salt bridge had been broken and all hydrogen bond interactions were removed, but surprisingly the stability of the structure increased although not significant. In a French study, this variant was found in 783 healthy individuals and none in AD patients but no experimental studies have been reported (Pottier et al., 2013). Finally, the D86V caused significant damage to TREM2 as a charged residue was replaced with an uncharged and removal of hydrogen bond interactions. The D86V variant was first found in a patient with FTD-like symptoms, which was also observed in a separate Turkish cohort (Guerreiro et al., 2013; Guerreiro et al., 2013). Experimental studies have shown that the substitution also results in impaired cell maturation and decreased cell surface expression (Sirkis et al., 2017).

Overall, it utilizes the AF2 structure of TREM2 to evaluate the stereochemical, functional, and structural effects of missense mutations that lead to the pathogenesis of AD, NHD, and FTD as well as those that remain unknown in current literature. Our data of NHD-causing variants support findings from prior studies, and provide additional information on the W50C mutation. For variants located on the surface of the IgV-like domain, we identified 10 that caused significant destabilizing effects to the receptor. Excluding variants implicated in NHD, we found that most of the mutations buried in the receptor did not cause significant changes to the overall protein, and further studies are required to explore their functional role. Ultimately, we believe that the in-silico biochemical data presented in this study for 32 variants will be informative to both clinical and experimental research in the near future.

## 5. Limitations

Although we provide novel biochemical findings for a majority of missense variants in TREM2, it must be noted that there are multiple limitations to be considered from this data. These limitations are directly related to our pilot study reported in the journal, revolving around the in-silico protocol (Pillai et al., 2025). In the original study, our protocol nearly accurately predicted all effects of the influential R47H variant, which we subsequently demonstrated on the Y38C, R62H, D87N and V126G variants that also validated prior literature. This inspired us to utilize this method for variants of unknown significance that can be evaluated on the AF2 structure. The key limitation of the in-silico mutagenesis techniques used to formulate the mutant structures is that they are based on rigid-structure predictions, and do not consider dynamic conformational changes. A notable instance is the R47H variant, where our predicted structure was unable to identify a hydrogen bond interaction with His67 (Pillai et al., 2025) although all other molecular details were correctly identified. We believe that the rigid-structure predictions are meant to serve as a reasonable baseline characterization but are currently prone to slight inaccuracies as shown on the experimental structures differing from AF2 in this study. Another limitation that must be recognized is the protein stability effects observed for all variants in this study. Due to the low sample size of computational tools, it is more challenging to ascertain the stability effects of all variants. We used the most rigorous statistical testing on the data collected, but a greater sample size is required in the future.

## Acknowledgements

J.P. acknowledges the OPALS program at the Institute of Engineering in Medicine, UC San Diego, USA for technical assistance.

## Author Contributions

J.P. and C.W. jointly conceived the study. All authors planned the entire experimental procedure presented. J.P. performed all experiments and analyzed the source data. All authors wrote the manuscript.

## Data Availability

All source data files are provided in the Supplementary Material and Supplementary Data. The source PDB files and PSE files made to formulate all figures have been provided at https://github.com/Joshua-Pillai/TREM2_Variants. The Prism file used for statistical analysis for this study has also been deposited in the GitHub repository.

## Supplementary Information

Supplementary Material - Details of variants of unknown significance and validation structures.

Supplementary Data - The source data for the structural and stability analyses. The 3D protein structures have been provided in the GitHub repository, including .PSE, .PDB, and .PRISM files.

---

**Fig S1.**
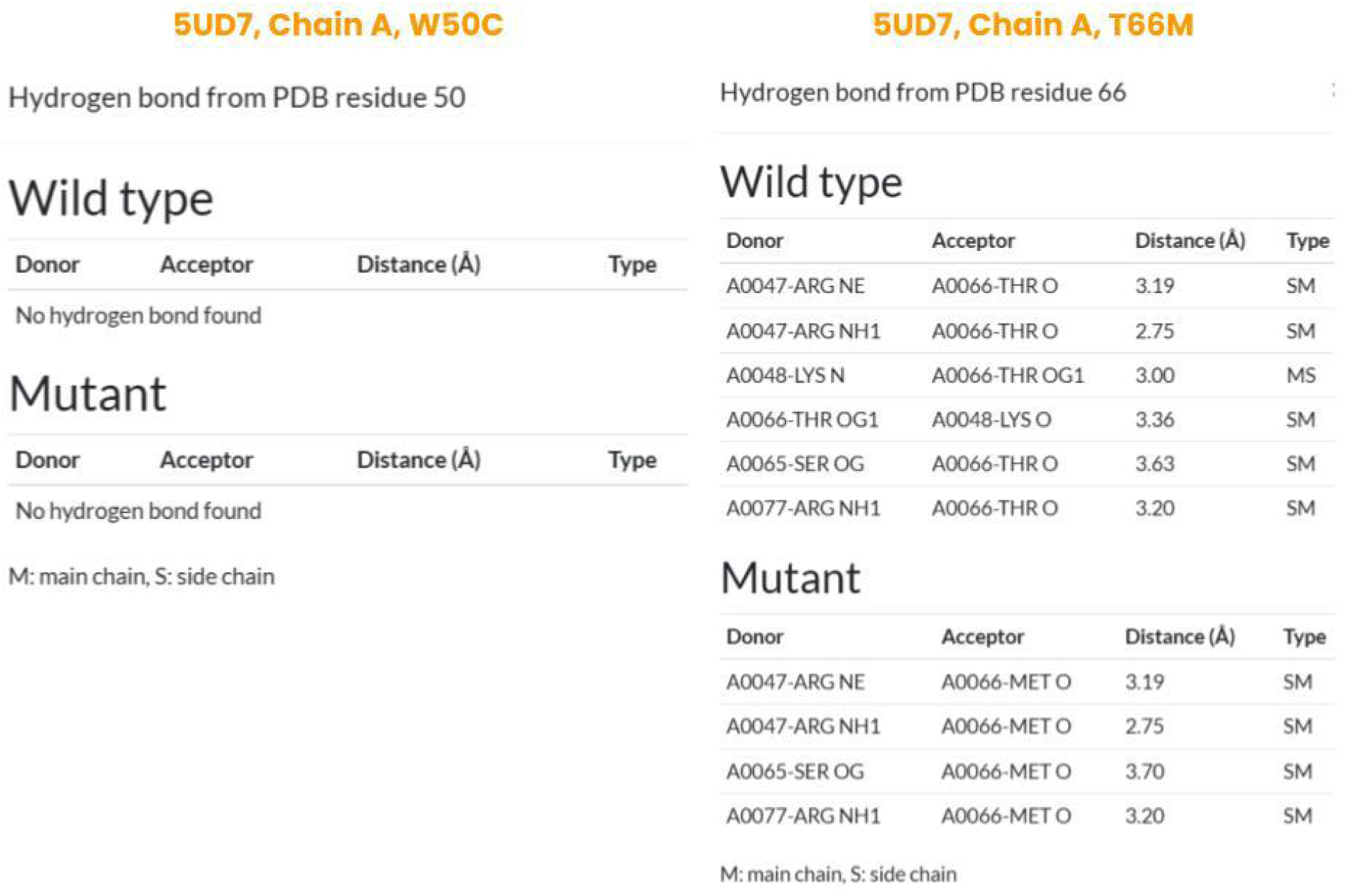

**Fig S2.**
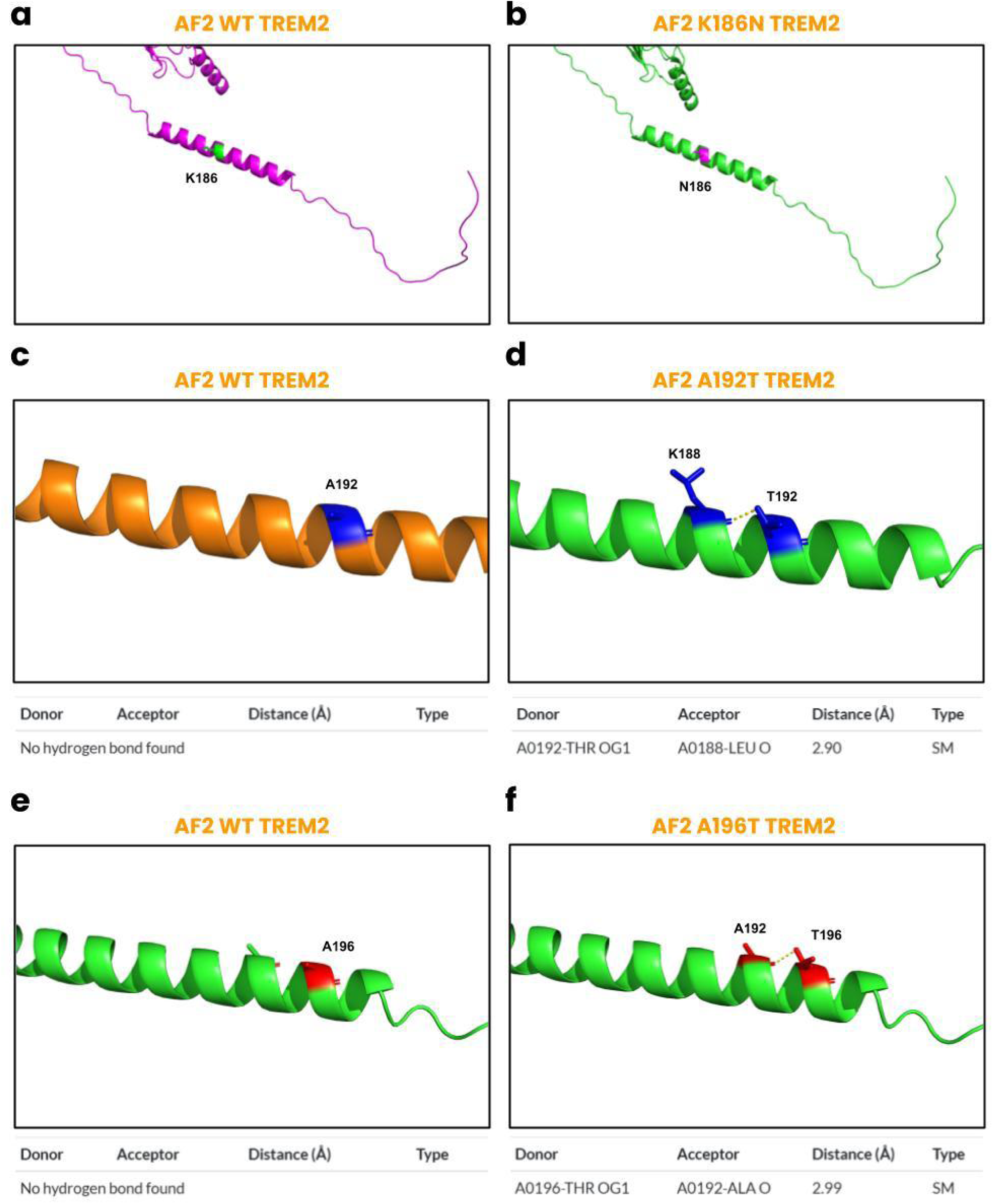

**Fig S3.**
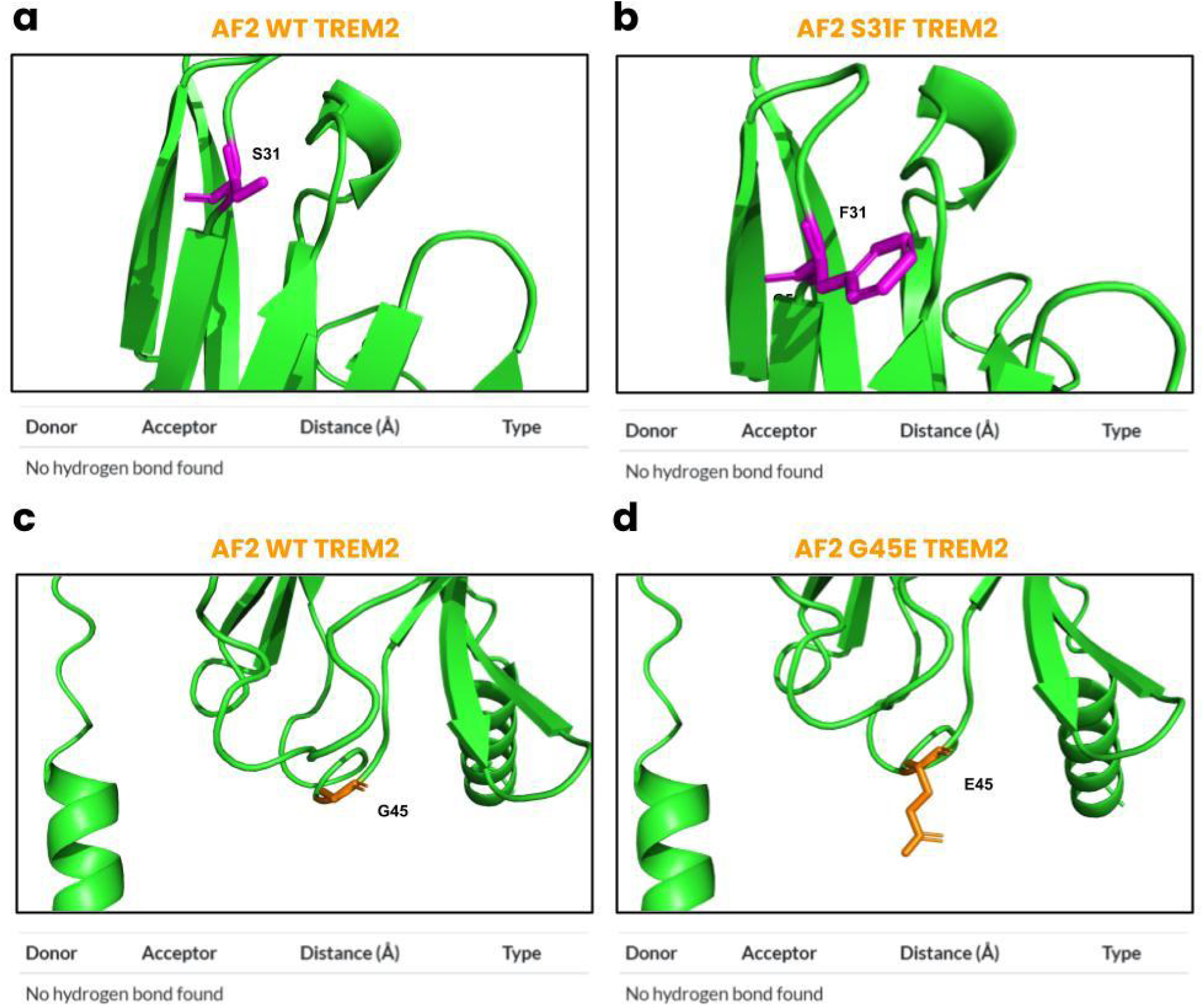

**Fig S4.**
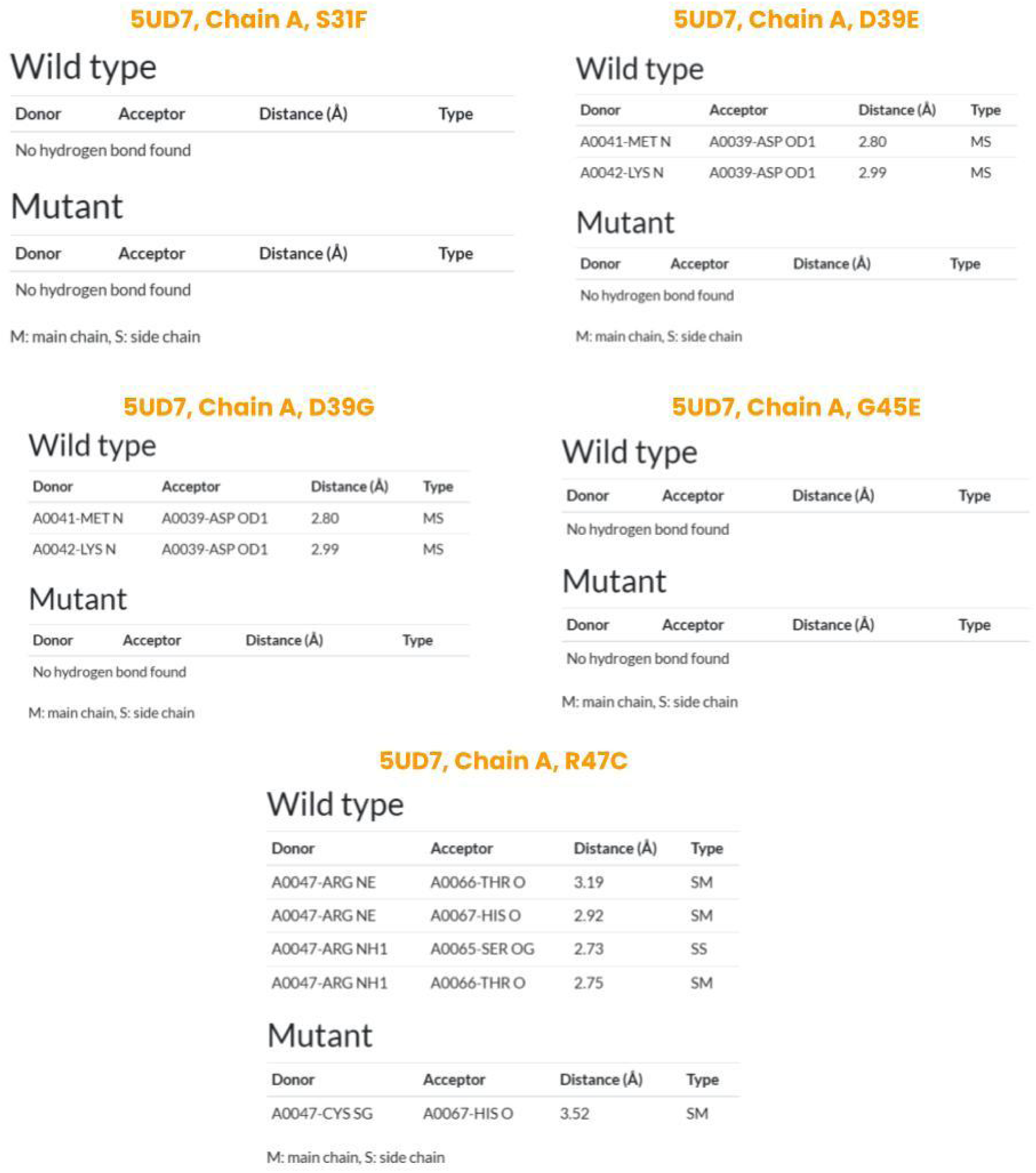

**Fig S5.**
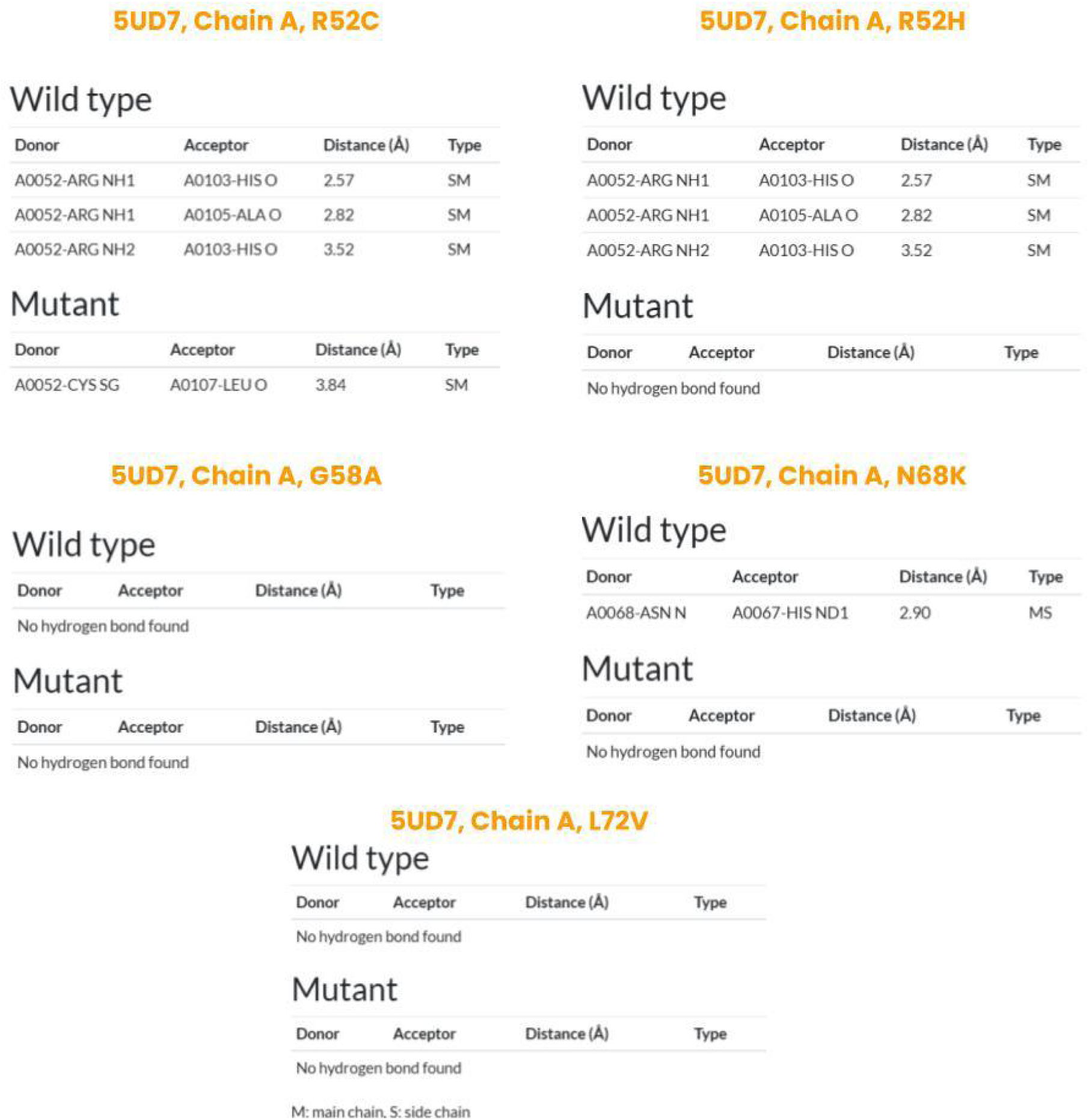

**Fig S6.**
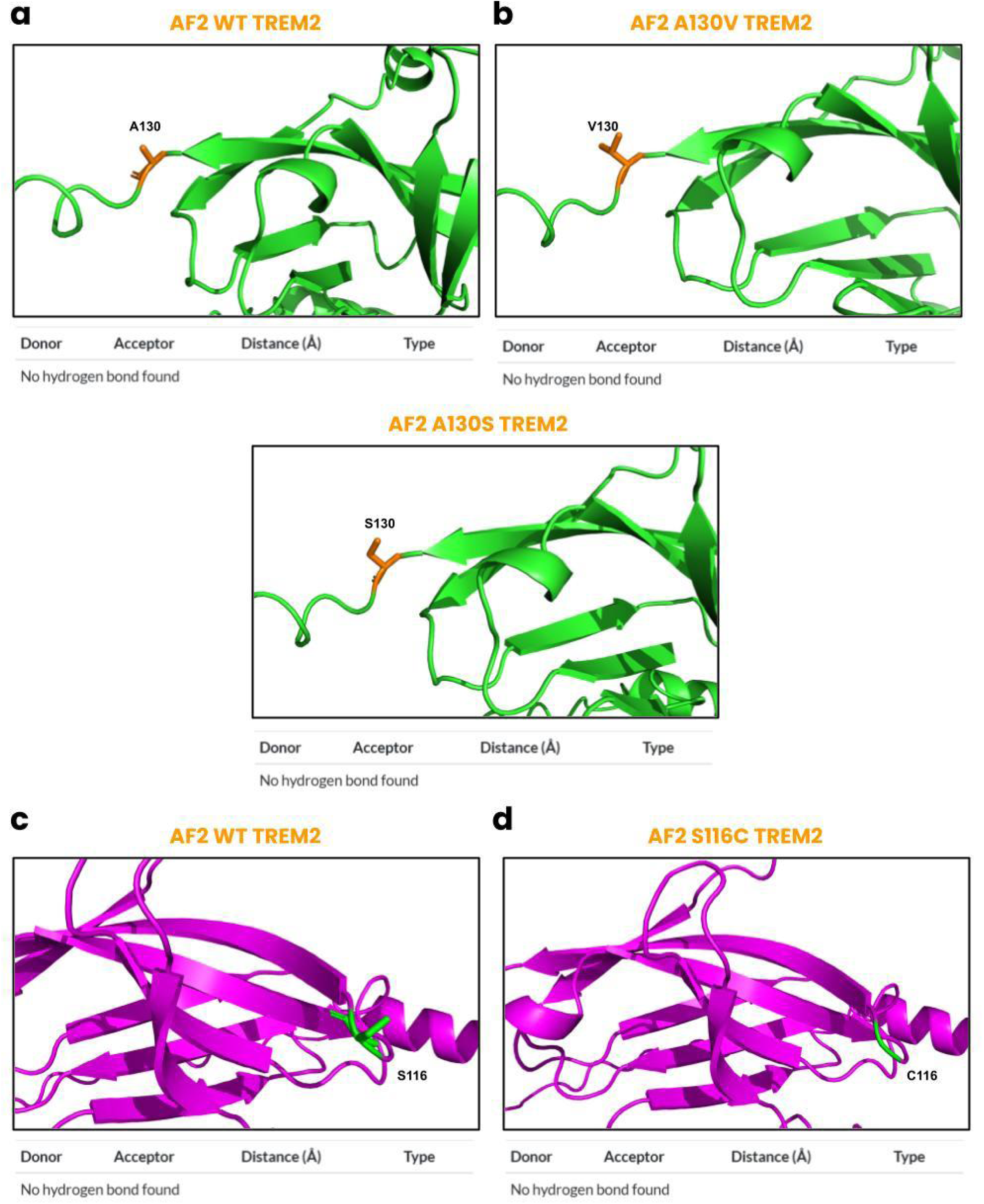

**Fig S7.**
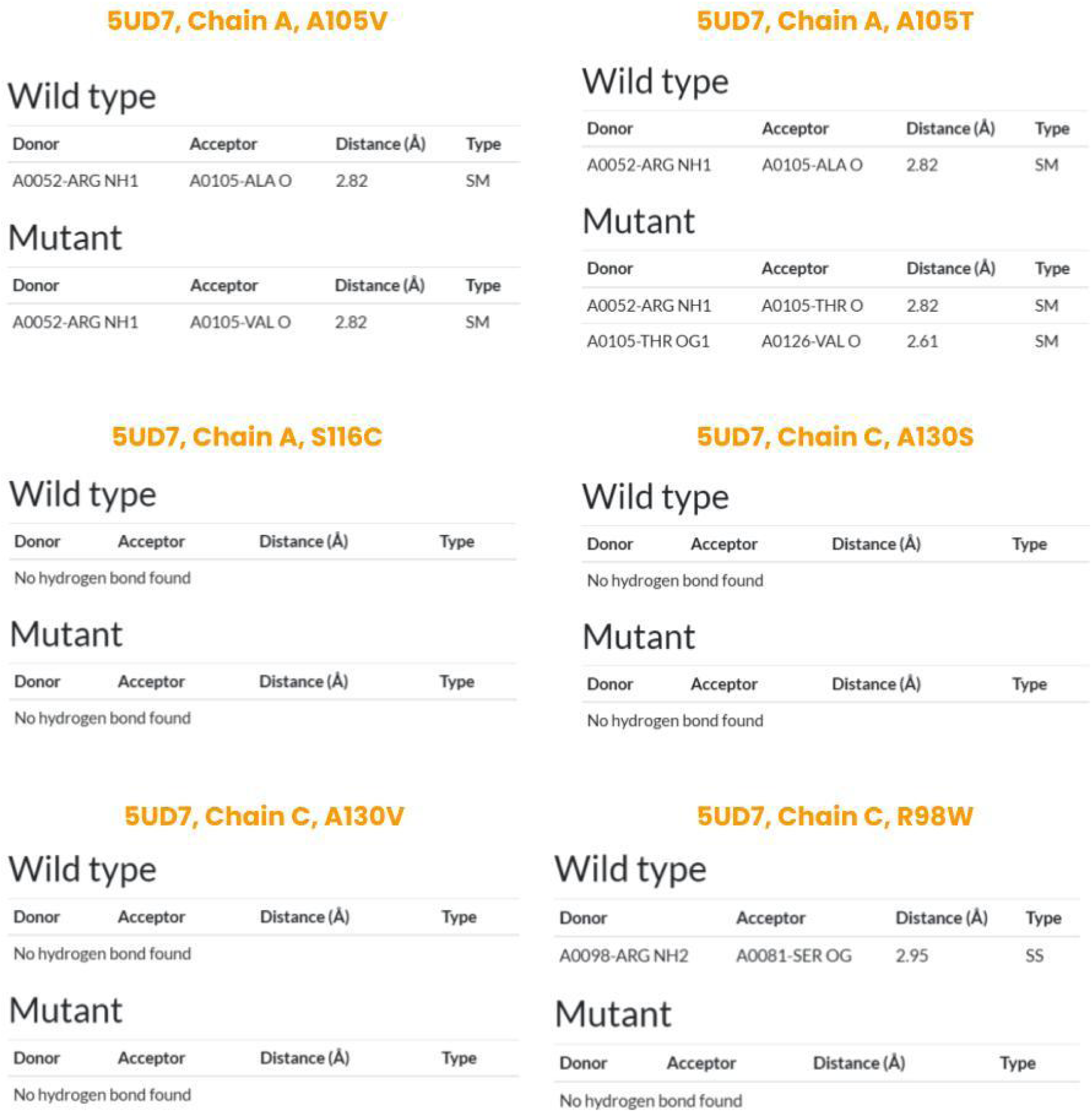

**Fig S8.**
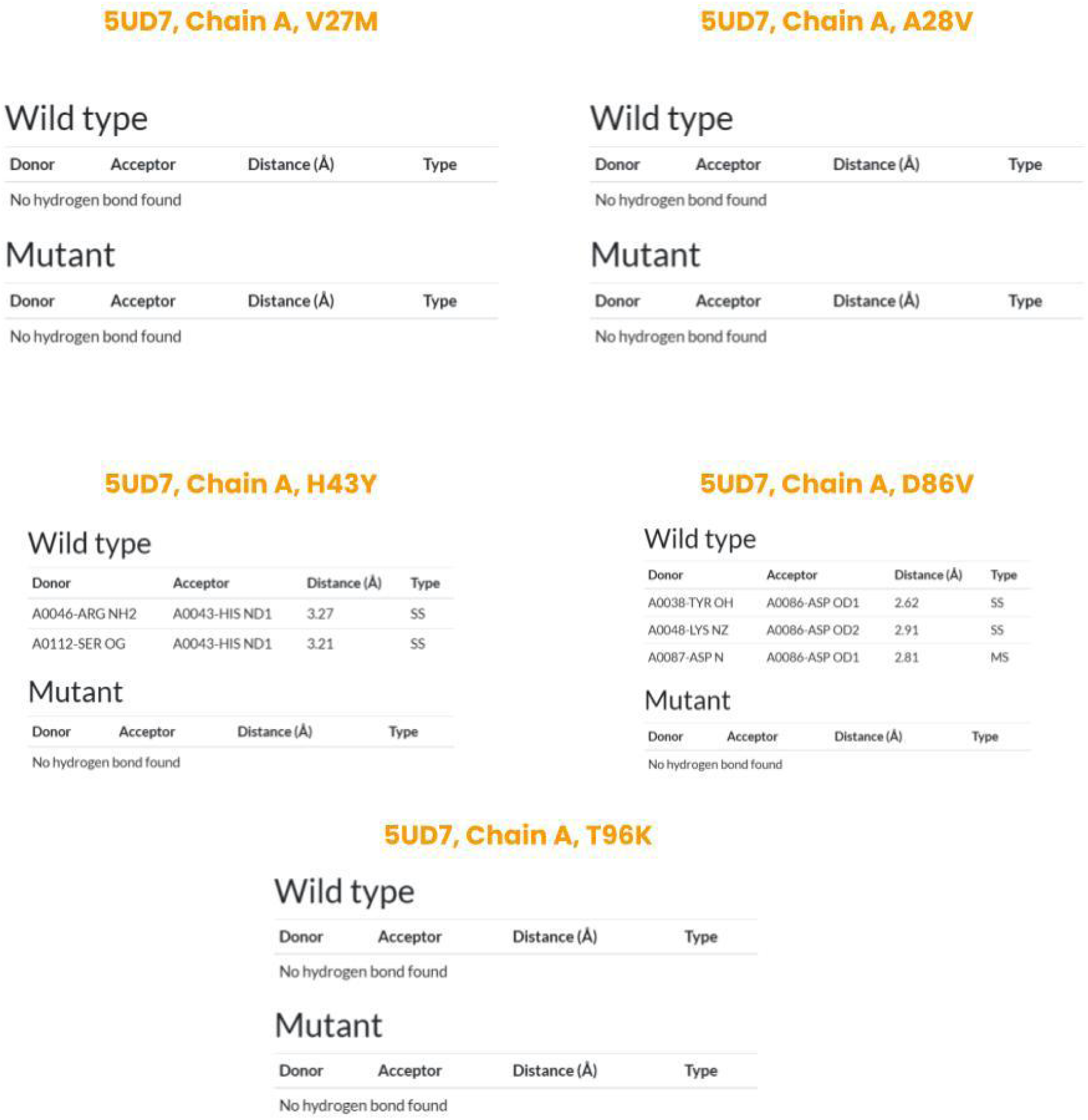

**Table S1.**
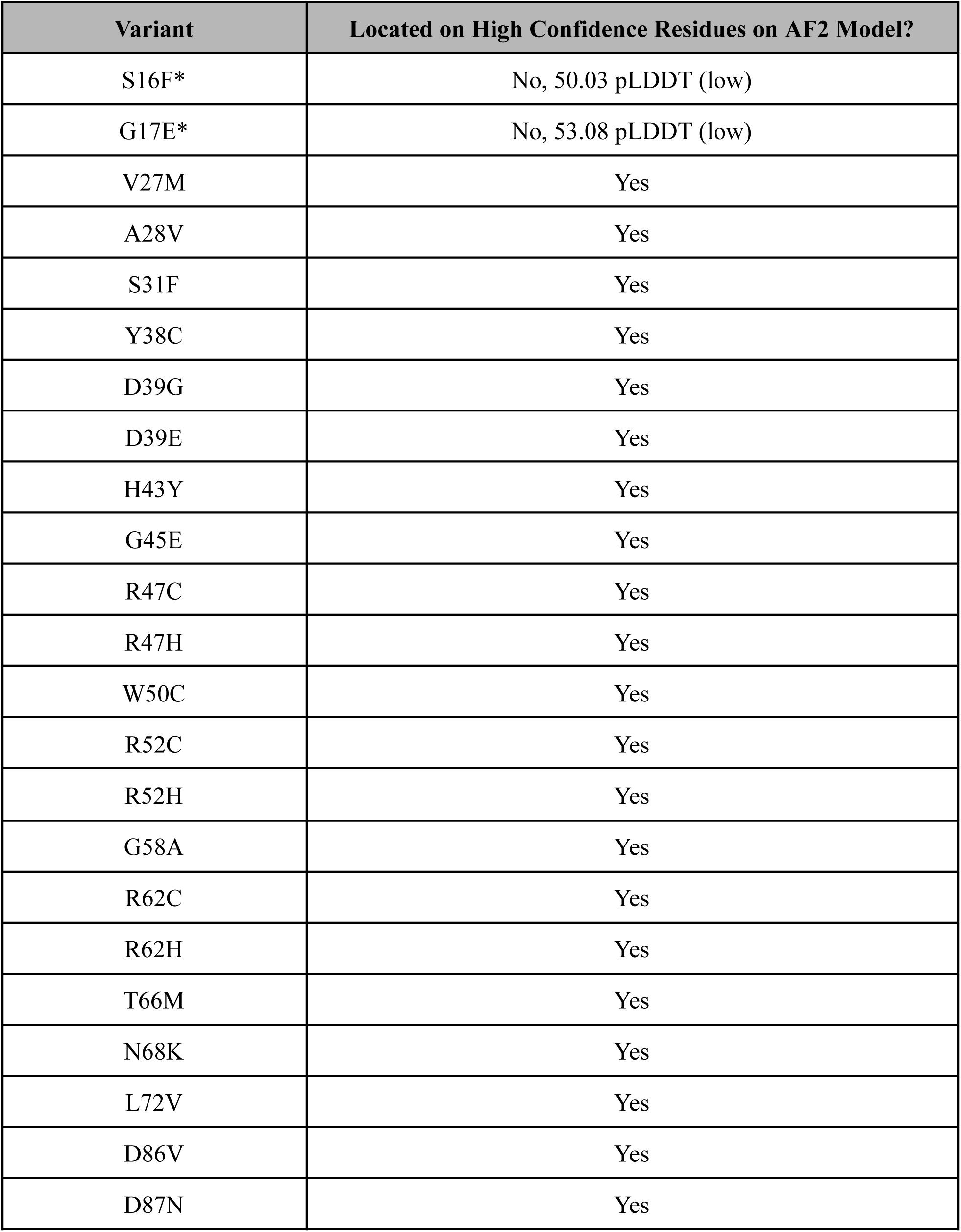
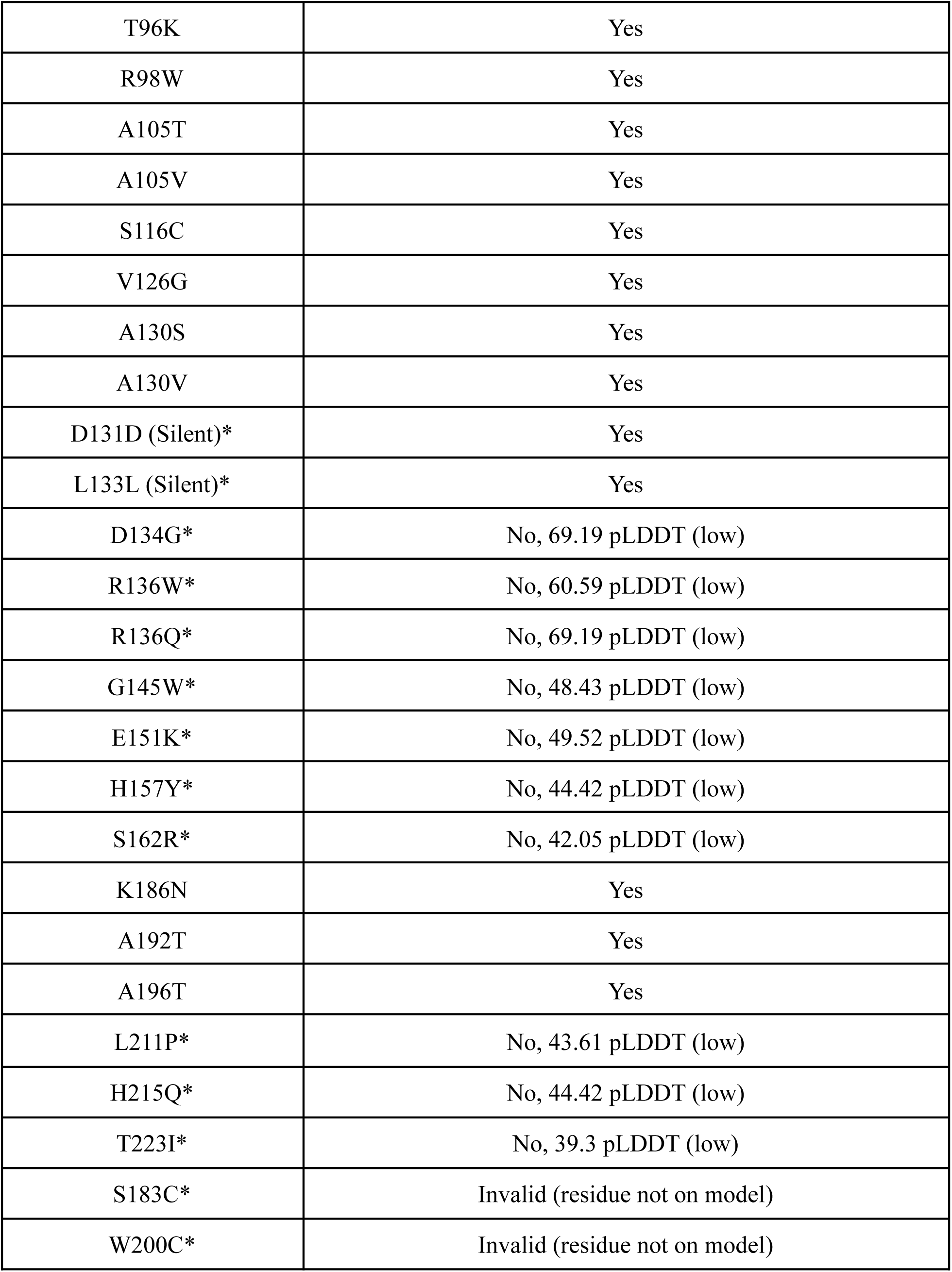
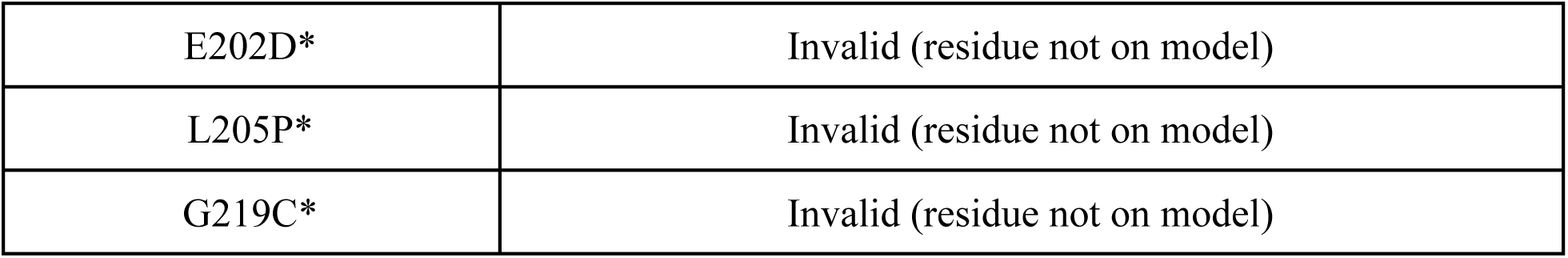
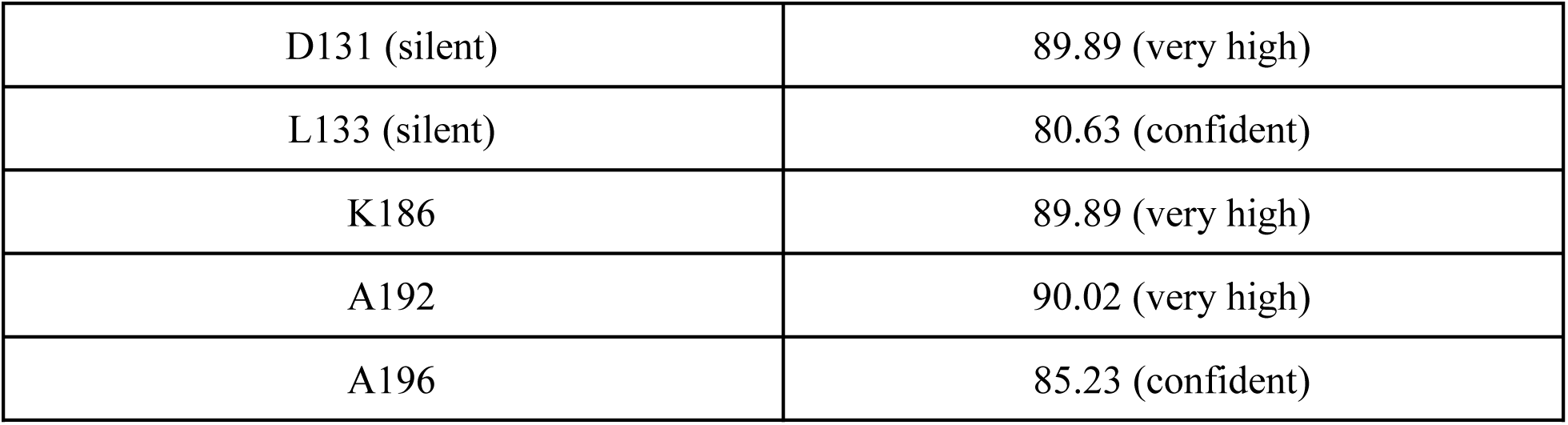
Missense mutations on *TREM2 (n = 51)*. Data presented in this table was obtained from Alzforum.org on January 31, 2025. 32 variants are located on high confidence residues.

**Table S2.**
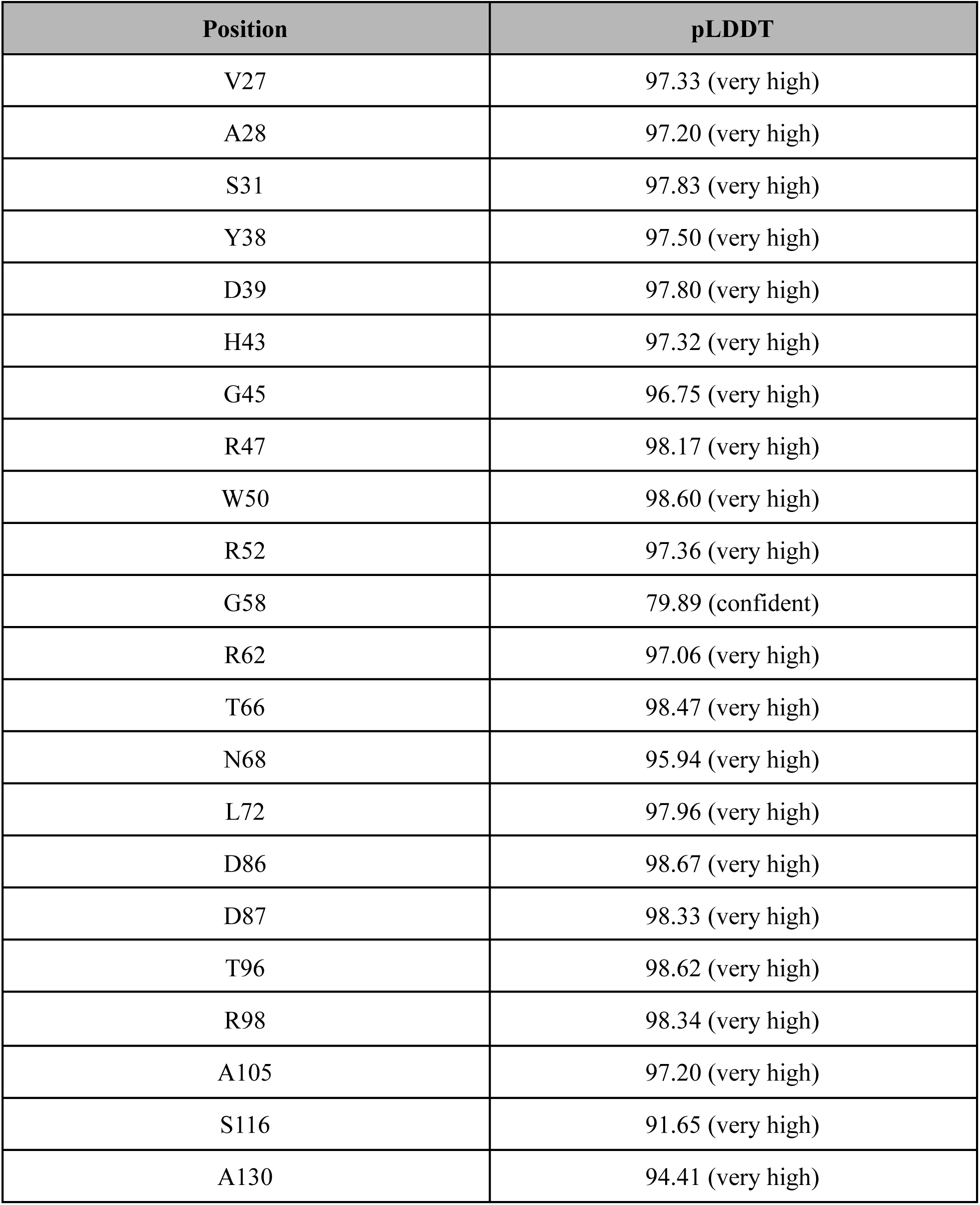
Predicted local distance difference test (pLDDT) score for amino acid positions of unknown variants in *TREM2*. Data presented in this table was obtained from the AlphaFold Structure Database on January 31, 2025.

**Table S3.**
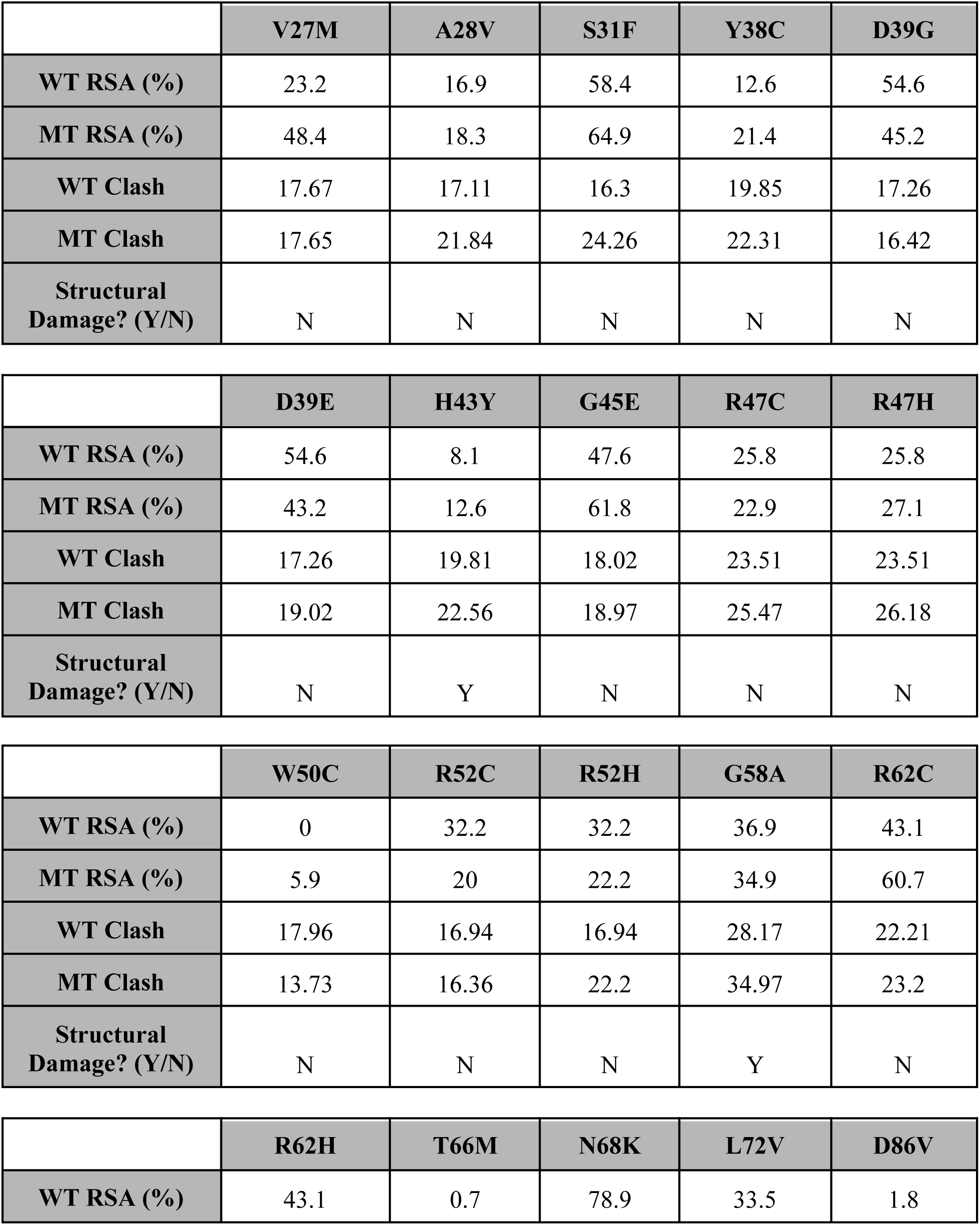
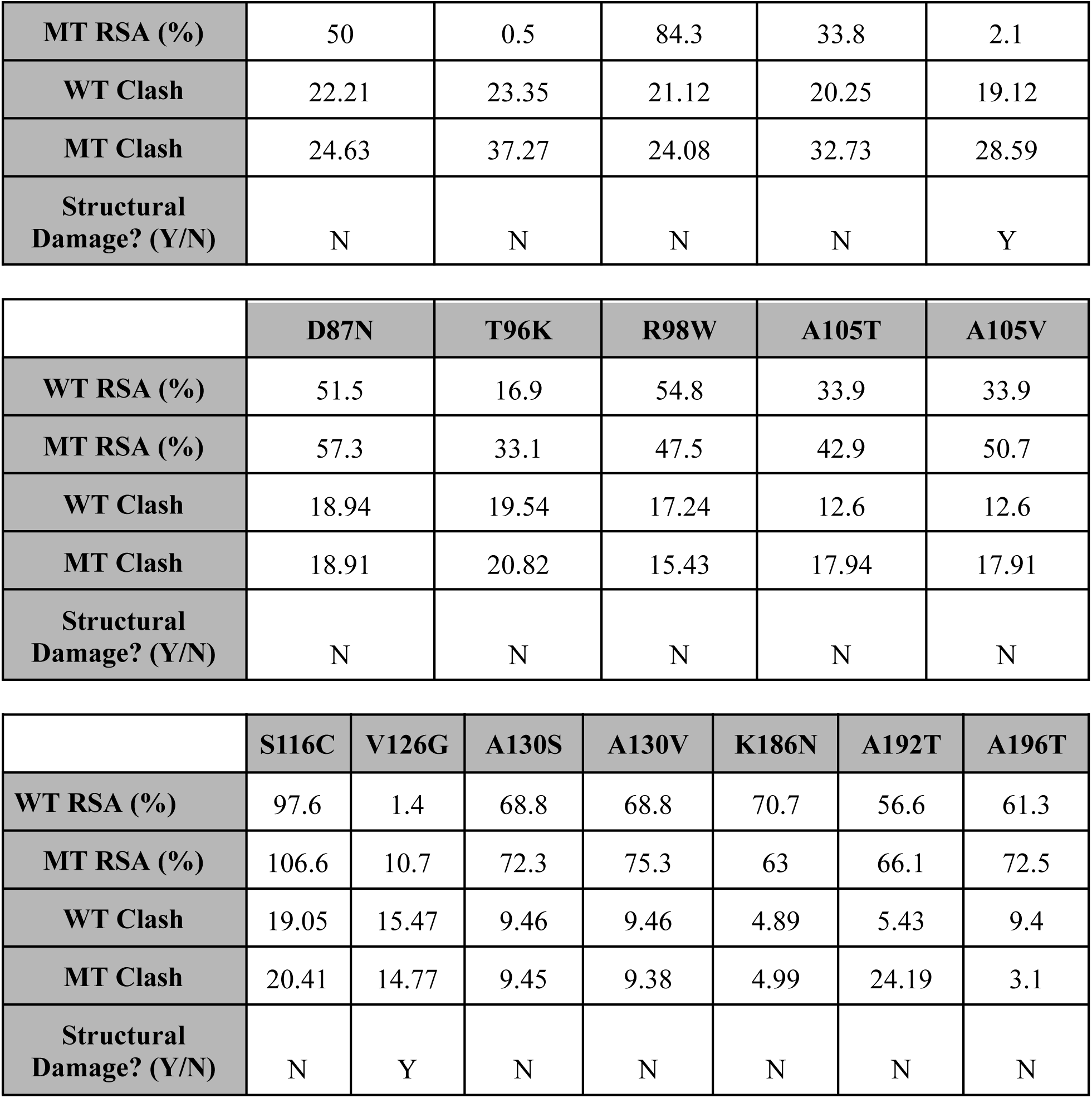
Source data of structural analyses of TREM2 variants. The data was collected on January 31, 2025.

|  | **V27M** | **A28V** | **S31F** | **Y38C** | **D39G** |
| --- | --- | --- | --- | --- | --- |
| **WT RSA (%)** | 23.2 | 16.9 | 58.4 | 12.6 | 54.6 |
| **MT RSA (%)** | 48.4 | 18.3 | 64.9 | 21.4 | 45.2 |
| **WT Clash** | 17.67 | 17.11 | 16.3 | 19.85 | 17.26 |
| **MT Clash** | 17.65 | 21.84 | 24.26 | 22.31 | 16.42 |
| **Structural Damage? (Y/N)** | N | N | N | N | N |

|  | **D39E** | **H43Y** | **G45E** | **R47C** | **R47H** |
| --- | --- | --- | --- | --- | --- |
| **WT RSA (%)** | 54.6 | 8.1 | 47.6 | 25.8 | 25.8 |
| **MT RSA (%)** | 43.2 | 12.6 | 61.8 | 22.9 | 27.1 |
| **WT Clash** | 17.26 | 19.81 | 18.02 | 23.51 | 23.51 |
| **MT Clash** | 19.02 | 22.56 | 18.97 | 25.47 | 26.18 |
| **Structural Damage? (Y/N)** | N | Y | N | N | N |

|  | **W50C** | **R52C** | **R52H** | **G58A** | **R62C** |
| --- | --- | --- | --- | --- | --- |
| **WT RSA (%)** | 0 | 32.2 | 32.2 | 36.9 | 43.1 |
| **MT RSA (%)** | 5.9 | 20 | 22.2 | 34.9 | 60.7 |
| **WT Clash** | 17.96 | 16.94 | 16.94 | 28.17 | 22.21 |
| **MT Clash** | 13.73 | 16.36 | 22.2 | 34.97 | 23.2 |
| **Structural Damage? (Y/N)** | N | N | N | Y | N |

|  | **R62H** | **T66M** | **N68K** | **L72V** | **D86V** |
| --- | --- | --- | --- | --- | --- |
| **WT RSA (%)** | 43.1 | 0.7 | 78.9 | 33.5 | 1.8 |
| **MT RSA (%)** | 50 | 0.5 | 84.3 | 33.8 | 2.1 |
| **WT Clash** | 22.21 | 23.35 | 21.12 | 20.25 | 19.12 |
| **MT Clash** | 24.63 | 37.27 | 24.08 | 32.73 | 28.59 |
| **Structural Damage? (Y/N)** | N | N | N | N | Y |
|  | **D87N** | **T96K** | **R98W** | **A105T** | **A105V** |
| **WT RSA (%)** | 51.5 | 16.9 | 54.8 | 33.9 | 33.9 |
| **MT RSA (%)** | 57.3 | 33.1 | 47.5 | 42.9 | 50.7 |
| **WT Clash** | 18.94 | 19.54 | 17.24 | 12.6 | 12.6 |
| **MT Clash** | 18.91 | 20.82 | 15.43 | 17.94 | 17.91 |
| **Structural Damage? (Y/N)** | N | N | N | N | N |

|  |  |  |  |  |  |  |  |
| --- | --- | --- | --- | --- | --- | --- | --- |
|  | **S116C** | **V126G** | **A130S** | **A130V** | **K186N** | **A192T** | **A196T** |
| **WT RSA (%)** | 97.6 | 1.4 | 68.8 | 68.8 | 70.7 | 56.6 | 61.3 |
| **MT RSA (%)** | 106.6 | 10.7 | 72.3 | 75.3 | 63 | 66.1 | 72.5 |
| **WT Clash** | 19.05 | 15.47 | 9.46 | 9.46 | 4.89 | 5.43 | 9.4 |
| **MT Clash** | 20.41 | 14.77 | 9.45 | 9.38 | 4.99 | 24.19 | 3.1 |
| **Structural Damage? (Y/N)** | N | Y | N | N | N | N | N |

**Table S4.** Source data of stability analyses of TREM2 variants. The data was collected on January 31, 2025.

|  | **V27M** | **A28V** | **S31F** | **Y38C** | **D39G** |
| --- | --- | --- | --- | --- | --- |
| **DynaMut2** | -0.22 | 0.71 | 0.26 | -1.37 | -0.12 |
| **DDMUT** | -0.26 | -0.5 | -0.7 | -1.82 | -0.53 |
| **PremPS** | 0.03 | 0.485 | -0.113 | -1.455 | -0.911 |
| **MAESTRO** | -0.42 | -0.55 | -0.02 | -1.93 | -1.24 |
| **DeepDDG** | -0.2364875 | 0.77650895 | -0.901268 | -1.838855 | -0.7633145 |
| **INPS-MD** | 0.234 | 0.636 | 0.365 | -0.683 | -0.845 |

|  | **D39E** | **H43Y** | **G45E** | **R47C** | **R47H** |
| --- | --- | --- | --- | --- | --- |
| **DynaMut2** | -0.26 | 0.68 | -0.21 | -0.53 | -0.57 |
| **DDMUT** | -0.06 | 0.85 | -0.5 | -0.71 | -0.99 |
| **PremPS** | -0.424 | 1.058 | -0.731 | -0.927 | -0.921 |
| **MAESTRO** | -0.41 | 0.46 | -0.06 | -0.04 | -0.77 |
| **DeepDDG** | -0.02171745 | 0.361917 | -0.131793 | -0.7192245 | -0.6663935 |
| **INPS-MD** | -0.27 | -1.192 | -0.19 | -0.848 | -0.843 |

|  | **W50C** | **R52C** | **R52H** | **G58A** | **R62C** |
| --- | --- | --- | --- | --- | --- |
| **DynaMut2** | -1.61 | -0.41 | -0.54 | 0.1 | -0.25 |
| **DDMUT** | -1.13 | -0.75 | -1.07 | -0.32 | -0.32 |
| **PremPS** | -2.639 | -0.525 | -0.425 | -0.771 | -0.273 |
| **MAESTRO** | -0.81 | 0.09 | -1.07 | -0.31 | 0.27 |
| **DeepDDG** | -1.58646 | -0.32159 | -0.767745 | -0.580008 | -0.34816 |
| **INPS-MD** | -1.717 | -0.112 | -0.7853600436 | -0.1881271 | -0.5693052 |

|  | **R62H** | **T66M** | **N68K** | **L72V** | **D86V** |
| --- | --- | --- | --- | --- | --- |
| **DynaMut2** | -0.38 | -0.91 | -0.02 | -0.24 | -0.28 |
| **DDMUT** | -1.03 | -0.52 | -0.11 | -0.86 | 0.07 |
| **PremPS** | -1.678 | -1.074 | 0.093 | -0.607 | 0.362 |
| **MAESTRO** | -0.068 | -1.2 | 0.1 | -0.71 | 0.15 |
| **DeepDDG** | 0.1471698 | -0.7927205 | 0.0675466 | -0.6506575 | -0.2352233 |
| **INPS-MD** | -0.504 | -0.8871422 | -0.1458005 | -0.9612144 | 0.9214804 |

|  | **D87N** | **T96K** | **R98W** | **A105T** | **A105V** |
| --- | --- | --- | --- | --- | --- |
| **DynaMut2** | 0.16 | -0.73 | -0.01 | 0.04 | -0.03 |
| **DDMUT** | 0.15 | -0.45 | -0.67 | -1 | -0.85 |
| **PremPS** | -0.337 | -1.533 | -0.072 | -0.501 | -0.507 |
| **MAESTRO** | -0.71 | 0.53 | 0.31 | -0.82 | -0.76 |
| **DeepDDG** | -0.0730616 | -1.308067 | -0.70065 | -0.48792735 | -1.146515 |
| **INPS-MD** | 0.043 | -1.021788 | 0.1985467136 | -0.1143406 | 0.089085313 |

|  | **S116C** | **V126G** | **A130S** | **A130V** | **K186N** | **A192T** | **A196T** |
| --- | --- | --- | --- | --- | --- | --- | --- |
| **DynaMut2** | -0.2 | -2.45 | -0.1 | 0.77 | 0.62 | 0.12 | 0.09 |
| **DDMUT** | -0.44 | -2.79 | 0.12 | -0.57 | 0.25 | 0.03 | 0 |
| **PremPS** | -0.095 | -1.678 | -0.155 | -0.055 | -0.738 | -0.061 | -0.167 |
| **MAESTRO** | 0.06 | -3.14 | -0.04 | -0.04 | -0.07 | 0.01 | 0.08 |
| **DeepDDG** | -0.3597165 | -3.952 | 0.0397 | 0.659496 | -0.6712 | -0.1810 | -0.542278 |
| **INPS-MD** | -0.1211262 | -2.898 | -0.2403222 | 0.08845126 | -0.084551 | -1.261405 | -1.065315 |

